# A perivascular niche supports endometrial epithelial regeneration

**DOI:** 10.1101/2024.03.07.583958

**Authors:** Shu-Yun Li, Sarah Whiteside, Bo Li, Xiaofei Sun, Tony DeFalco

## Abstract

Endometrial regeneration is essential for reproductive cycles and pregnancies, allowing the endometrium to undergo repair and renewal after menstruation and parturition. Epithelial cells lining the uterine cavity are shed during each cycle, although remnant luminal and glandular epithelial cells can regenerate the lumen lining. It is presumed that adult stem/progenitor cells in the uterine stroma also contribute to this regeneration. However, the specific cell type(s) and the underlying mechanisms have not been determined. Herein, we use genetic lineage tracing assays in mice to identify *Nestin*+ perivascular cells as active contributors to epithelial regeneration. Notch signaling maintains *Nestin*+ perivascular cells in a quiescent state but these cells re-enter the cell cycle and differentiate into epithelial cells via estrogen-stimulated suppression of Notch signaling dependent on estrogen receptor alpha (ERα/ESR1). These findings demonstrate that perivascular cells support epithelial regeneration and reveal a mechanism regulating the quiescence and activation of uterine perivascular cells.

## Introduction

The endometrium, composed of epithelial (glandular and luminal) and stromal compartments, is the uterine lining essential for the proper establishment and maintenance of a successful pregnancy ^1,2^. It is a highly dynamic tissue, undergoing morphological and functional changes in response to fluctuating sex steroid hormones during pregnancy and menstruation^3-5^. In mice, endometrial epithelial cell numbers increase approximately nine-fold from diestrus to estrus stages^6^. After the loss of a significant amount of epithelial cells during menses and pregnancy, the uterus must restore epithelial integrity to support future pregnancies^3-5,7,8^, and it is presumed that adult stem/progenitor cells in the remnant uterine tissue are responsible for its remarkable regenerative capacity^3,9^. The endometrial stromal bed contains various mesenchymal cells, including stromal fibroblasts, vascular/perivascular cells, and immune cells. Mesenchymal cells have been proposed as a source of epithelial progenitors^9,10^. Disorders of endometrial regeneration are closely related to severe pathological conditions, including infertility, endometriosis, endometrial hyperplasia, adenomyosis, and endometrial cancer^1,11-13^. However, the existence and exact identity of the epithelial progenitors remains elusive due to the heterogeneity of mesenchymal cells and the innate regenerative ability of epithelia.

The rapid restoration of endometrial integrity is thought to be due at least in part to the mobilization and differentiation of tissue-resident stromal mesenchymal stem cells^9^. Previous studies have described cells transitioning between a stromal and epithelial identity that are recruited to breaches in the uterine lining, suggesting that mesenchymal cells are endometrial epithelial progenitors^9,10^. The role of stromal cells has been further investigated in mouse models of menstruation and postpartum repair, using *Amhr2*-Cre transgenic mice to lineage trace mesenchymal cells and their progeny. These studies suggest that *Amhr2*+ cells partly contribute to endometrial re-epithelialization after menses and parturition^14,15^. Using a *Pdgfrb*-BAC-eGFP knock-in mouse model, in which GFP expression is restricted to endometrial mesenchymal cells, GFP was expressed in actively repairing regions where the stromal surface is denuded after decidual shedding and a new epithelial cell layer is being re-established^16^. In particular, perivascular cells are proposed to be stem cells within the endometrial stroma because they have some common phenotypes and share gene expression patterns with mesenchymal stem cells^9,17^. A recent single-cell analysis of human endometrial perivascular cells revealed that both secretory and menstrual perivascular cells expressed mesenchymal-stem-cell-specific genes once they leave the perivascular niche, indicating a high potential of differentiation. However, due to the lack of an ability to trace perivascular cells in human endometrium, it is unknown whether they are involved in human endometrial regeneration^18^.

Notch signaling is a highly conserved multifunctional cell communication system that plays a crucial role in various biological processes, including embryonic development, tissue homeostasis, and cell fate determination. Notch receptors and ligands are transmembrane proteins that interact through juxtacrine signaling, initiating a cascade of molecular events that regulate cellular differentiation, proliferation, and apoptosis^19,20^. Notch signaling has been implicated in regulating the uterus as it undergoes remarkable changes during the menstrual cycle, pregnancy, and postpartum period, involving intricate interactions between various cell types to enable successful implantation, decidualization, and embryo development^3-5,21-24^. Activation of Notch signaling by Jagged1 (JAG1) is required for the maintenance of human endometrial mesenchymal stromal/stem-like cells *in vitro*^25,26^.

Nestin is a marker of multiple stem/progenitor cell types, including neural stem cells and those in the hematopoietic system^27,28^. Recent single-cell transcriptome studies of endometrium suggested that both human and murine perivascular cells highly express *Nestin*^16,18^. It has been challenging to create a reporter mouse line that faithfully and specifically replicates the endogenous expression pattern of perivascular cells. Multiple *Nestin*-CreER mouse lines have been created to trace the cell fate of *Nestin*+ cells^29-34^. Recently we reported that one^34^ of these *Nestin*-CreER mouse lines effectively recapitulated endogenous *Nestin* expression in the gonad, and we demonstrated that *Nestin*+ perivascular cells are multipotent progenitors in the fetal testis and ovary^35,36^. Here, using the same *Nestin*-CreER mice, we observed that Nestin is expressed in endometrial perivascular cells and exhibits a distinct expression pattern as compared to NG2, another pericyte marker (official name CSPG4), during endometrial regeneration. We employed genetic lineage tracing assays to explore the plasticity of perivascular cells in endometrial regeneration using long-term lineage tracing, pregnancy, and artificial decidualization mouse models. Intriguingly, our analyses revealed that these *Nestin*+ perivascular cells contribute to epithelial expansion and regeneration during estrous cycles and endometrial repair. Endometrial regeneration is an estrogen-dependent process^3^, and an ovariectomized mouse model showed that estrogen induces the differentiation of *Nestin*+ perivascular cells in an estrogen receptor alpha (ERα/ESR1)-dependent manner. Using a Notch signaling reporter mouse strain, we found that Notch activity, likely through DLL4/Notch3 interactions, is enriched in perivascular cells. Conditional Notch loss-of-function in perivascular cells induced their exit from quiescence to an active state, indicating that Notch activity is required for maintaining quiescence of perivascular cells. Interestingly, Notch-mediated maintenance of quiescent status is overridden by estrogen signaling, and perivascular cells are activated upon exposure to estrogen. Thus, our findings reveal a novel role for perivascular cells in endometrial regeneration in an estrogen-dependent mechanism. Understanding the involvement of perivascular cells in endometrial regeneration holds considerable therapeutic potential for treating uterine-related disorders and improving women’s reproductive health.

## Results

### A *Nestin*-CreER mouse line faithfully marks uterine Nestin+ perivascular cells

To investigate the expression pattern of Nestin in the uterus, we first examined P60 (postnatal day 60) uteri using immunofluorescence analyses. We observed that Nestin protein was specifically localized in cells adjacent to endothelial cells, suggesting Nestin was expressed in uterine perivascular cells (Fig. 1A). To further investigate whether Nestin+ cells were perivascular cells, we used tamoxifen-inducible *Nestin-* CreER mice to permanently label *Nestin*+ cells and their progeny. To induce CreER activity, *Nestin*-CreER; *Rosa*-Tomato mice were exposed to 4-hydroxytamoxifen (4-OHT) at P60 and P61, and uteri were collected at P62. Perivascular *Nestin*+ cells were successfully labeled with Tomato in uteri (Fig. 1B). Our analyses revealed that around 50% of *Nestin*+ cells were labeled with Tomato (Fig. 1E). To further confirm the identity of Tomato+ perivascular cells, we examined an endothelial nuclear marker, ERG, and another pericyte marker, NG2. All Tomato+ cells were located next to ERG+ cells (but did not co-express ERG), and they co-expressed NG2 (Fig. 1C, D and F). Furthermore, no Tomato+ cells were detectable within CK8+ (cytokeratin 8+) epithelia (Fig. 1C, D and G). These data show that we successfully established a mouse model to label and track *Nestin*+ perivascular cells in the adult uterus.

**Figure 1.**
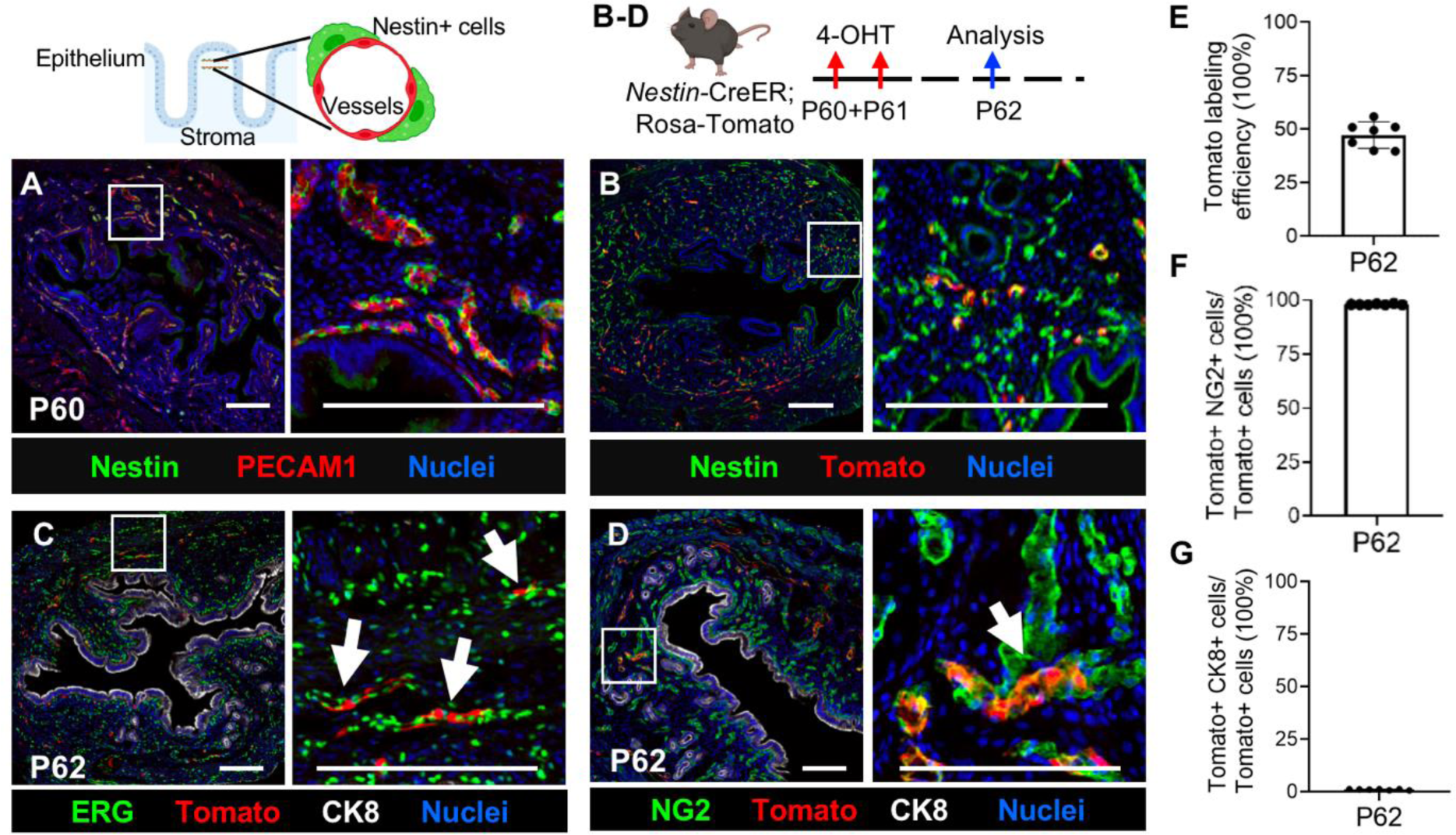
*Nestin*+ cells are endometrial perivascular cells. (A) Immunofluorescence images of P60 wild-type uterus. (B-D) Immunofluorescence images of P62 *Nestin*-CreER; *Rosa*-Tomato uteri exposed to 4-OHT at P60 and P61. Tomato (arrows) was not expressed in ERG+ endothelial cells (C) but was expressed in NG2+ perivascular cells (D). Scale bar: 200 μm. (E-G) Quantification of percent Nestin+ cells expressing Tomato (E), percent Tomato+ cells expressing NG2 (F), and percent Tomato+ cells expressing CK8 (epithelial marker) (G).

### Nestin+ perivascular cells contribute to epithelia during estrous cycles

The mouse endometrium undergoes significant epithelial turnover during estrous cycles^37^. To determine the fate of Tomato+ perivascular cells and their contribution to epithelia during cyclical endometrial turnover, Tomato+ cells were tracked for extended periods, spanning 1 month, 2 months, and 4 months post-injection. Our analyses showed that stromal Tomato+ cells persist throughout all time points examined (Fig. 2A, C, E, and G). We observed that the percentage of Tomato+ cells throughout the stroma (excluding the myometrium) peaked two months after 4-OHT induction at 12%, and then decreased to ∼4% at four months after induction (Fig. 2G). In all samples examined, the majority (∼60- 70%) of Tomato+ cells maintained NG2 expression and were localized adjacent to blood vessels (Fig. 2A, C and E; quantified in Fig. 2H). These results suggest that Tomato+ Nestin-derived cells are a long-lasting cell population localized adjacent to endothelial cells and retain perivascular cell markers.

**Figure 2.**
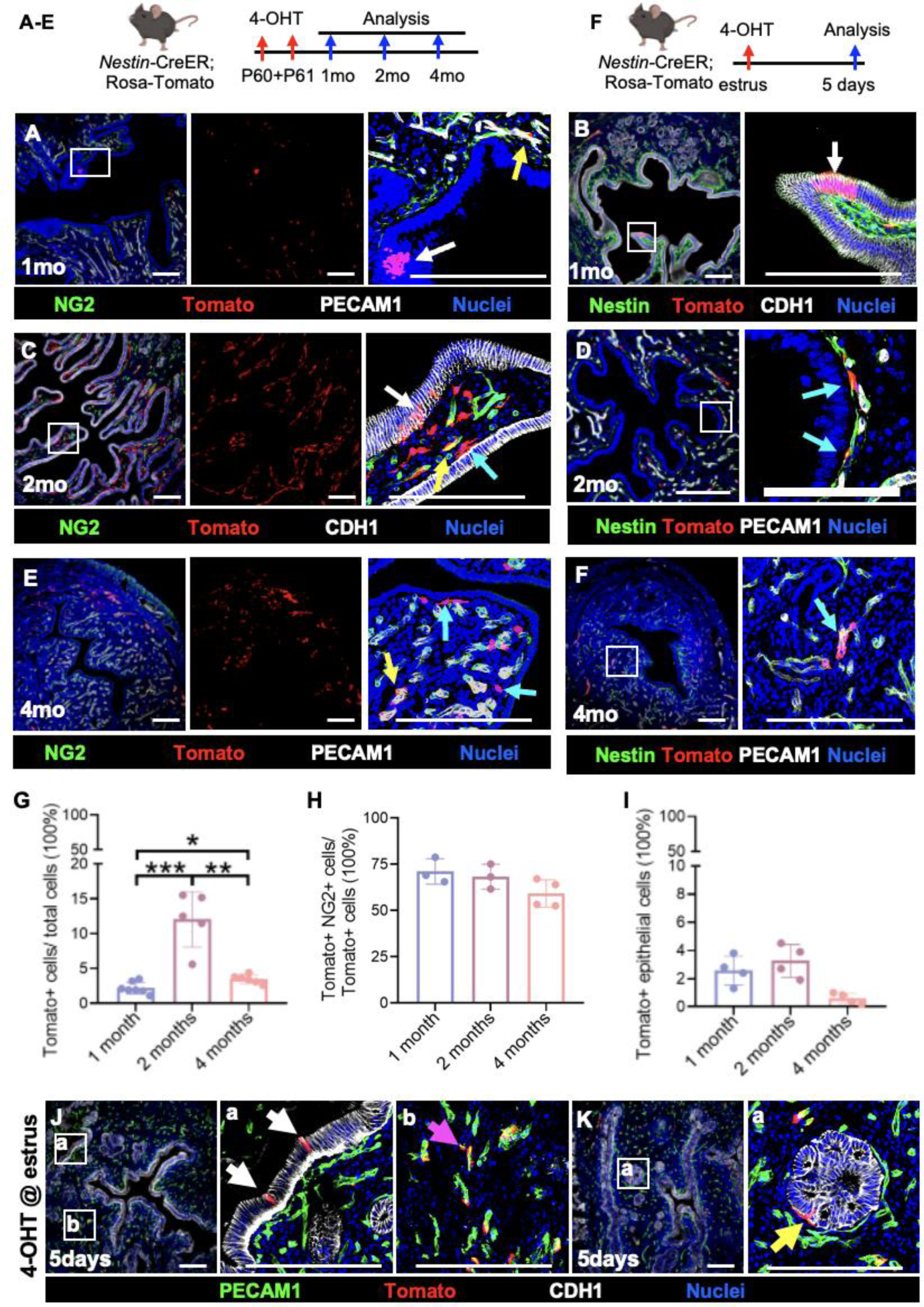
*Nestin*+ perivascular cells differentiate into epithelial cells during estrous cycles. (A-F) Long-term lineage tracing of *Nestin*+ perivascular cells in *Nestin*-CreER; *Rosa*-Tomato uteri 1 month (A, B), 2 months (C, D), and 4 months (E, F) after exposure to 4-OHT at P60 and P61. White arrows indicate Tomato-expressing cells in the epithelium (CDH1+); yellow arrows indicate Tomato-expressing cells that are NG2+; and cyan arrows point to Tomato-expressing cells that do not express NG2 or CDH1 (C and E), but still express Nestin (D and F). (G) Quantification of percent Tomato+ cells among all uterine cells (excluding myometrium). (H) Quantification of percent Tomato+ NG2+ cells among all Tomato+ cells. (I) Quantification of percent Tomato+ CDH1+ cells among all CDH1+ epithelial cells. (J and K) Short-term lineage tracing of *Nestin*+ perivascular cells in *Nestin*-CreER; *Rosa*-Tomato uteri 5 days after 4-OHT exposure during the estrus phase. White arrows indicate Tomato-expressing cells in the epithelium (CDH1+); magenta arrow indicates Tomato-expressing cells that are adjacent to PECAM1+ vasculature; and yellow arrow points to Tomato+ cell half-embedded in the epithelial layer. mo: month. Thin scale bar: 200 μm; thick scale bar: 100 μm. \**P*<0.05, \*\*\**P*<0.001.

A subset of Tomato+ cells differentiated into epithelial cells (CDH1+) in uteri 1 month and 2 months post-induction, while sporadic Tomato+ epithelial cells were observed in 4-month post-induction samples (Fig. 2A-G and I). In addition to Tomato+ cells that had incorporated into epithelia, additional Tomato+ cells were observed subjacent to epithelial cells. Tomato+ cells underneath epithelia retained Nestin, but not NG2, expression (Fig. 2C-F), in contrast to the observation that almost all perivascular Tomato+ cells expressed NG2 (see Fig. 1D and F). The results suggest perivascular Nestin+ cells migrate towards the epithelial layer with a gradual loss of NG2 expression, and eventually some of them incorporate into the epithelial layer. These data suggest the presence of a unique subpopulation of pericytes located adjacent to the uterine epithelium that selectively expresses Nestin, thus indicating that there is pericyte heterogeneity in the uterus.

Considering that both Nestin and NG2 displayed a pericyte-like expression pattern in non-pregnant uteri, we next examined uteri during early pregnancy to determine if hormonal changes affect their expression pattern. Nestin and NG2 exhibited a similar expression pattern in day 4 (D4) uteri, with both being expressed in cells adjacent to blood vessels (Fig. S1A). Interestingly, we observed a different expression pattern between Nestin and NG2 in D8 uteri (Fig. S1B). Nestin was predominantly expressed near blood vessels in the mesometrium, while NG2 was mainly expressed in the stroma of the anti-mesometrium. This suggests a dynamic shift in the spatial distribution of NG2 as pregnancy progresses, while Nestin was consistently expressed in cells adjacent to blood vessels. These data suggest that Nestin marks a persistent uterine perivascular cell population which is not influenced by pregnancy.

A previously published single-cell RNA-seq analysis revealed heterogeneity of *Nestin* and *Ng2* (*Cspg4*) expression in perivascular cells during endometrial regeneration^16^. We re-analyzed that published dataset, which focused on purified stromal cells from an artificial decidualization model, by re-clustering perivascular cell populations into four distinct subsets (Fig. S1C). The feature plots of *Ng2* and *Nestin* revealed that some perivascular cells are positive for only one of the two markers (Fig. S1D). A further quantification of percentages of *Ng2*+ and/or *Nestin*+ cells showed that perivascular cells are heterogeneous in *Ng2* and *Nestin* expression (Fig. S1E).

To examine whether Tomato-labeled pericytes participated in short-term epithelial expansion during the estrous cycle, a single dose of 4-OHT was administered to *Nestin*-CreER; *Rosa*-Tomato females during the estrus phase. We began detecting Tomato+ cells in the epithelium 5 days after injection (Fig. 2J and K). Interestingly, we also observed a Tomato+ cell half-embedded in the epithelial layer, suggesting it was undergoing a transition. In sum, most Tomato+ cells were maintained as perivascular cells, but a few Tomato+ cells contributed to epithelial expansion in endometrium during estrous cycles, suggesting that *Nestin*+ perivascular cells have the abilities of self-renewal and differentiation over short time scales.

### *Nestin*+ perivascular cells participate in epithelial regeneration during pregnancy

The mouse endometrium undergoes greater remodeling and regeneration during pregnancy as compared to estrous cycles^15^, since the loss of epithelial cells in implantation sites subsequently requires vast epithelial regeneration during pregnancy. We assessed Tomato expression in D12 *Nestin*-CreER; *Rosa*-Tomato females (exposed to 4-OHT at P60 and P61, prior to mating), when the epithelium is regenerating, and postpartum day (PPD) 3, when regeneration of epithelium is almost complete^14^. Tomato was observed in the newly regenerated epithelium in D12 and PPD3 endometrium (Fig. S2A and B). These data indicate that *Nestin*+ cells participate in epithelial regeneration during pregnancy. Given that the number of Tomato-labeled cells within the stroma peak at 2 months after 4-OHT injections in nulliparous mice (see Fig. 2G), we next explored the contribution of perivascular cells in uterine regeneration of postpartum *Nestin*-CreER; *Rosa*-Tomato females 2 months after exposure to 4-OHT. On PPD3, about 8% of epithelial cells were positive for Tomato (Fig. 3A and B). Furthermore, the percentage of Tomato+ epithelial cells in PPD3 uterus was significantly higher at 2 months post-induction than at 2 days post-induction (Fig. 3B).

**Figure 3.**
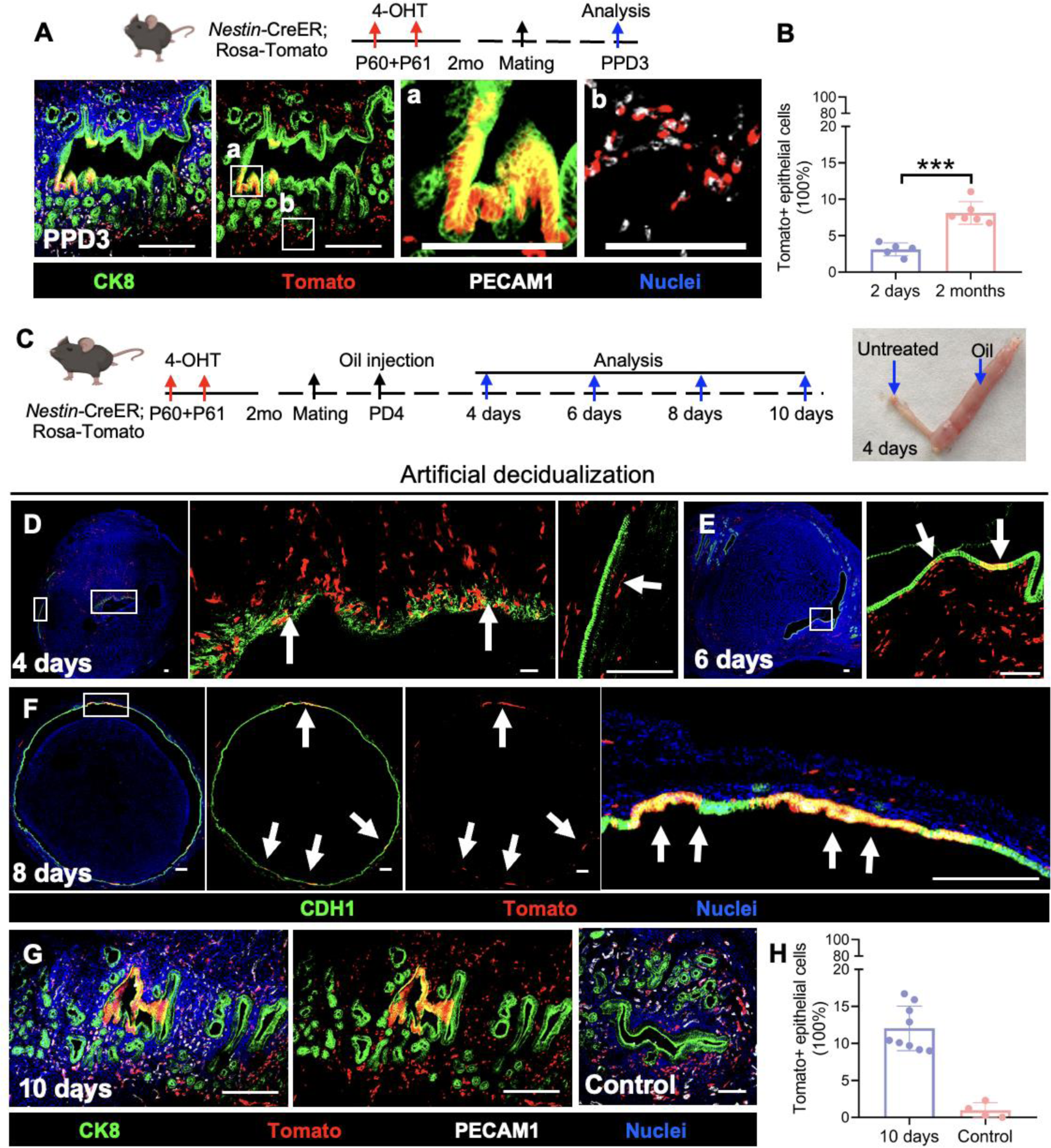
*Nestin*+ perivascular cells contribute to epithelial regeneration during endometrial regeneration. (A) Immunofluorescence images of *Nestin*-CreER; *Rosa*-Tomato PPD3 uteri 2 months after exposure to 4-OHT at P60 and P61. (B) Quantification of percent Tomato-positive epithelial cells at PPD3 either 2 days or 2 months after exposure to 4-OHT. (C) Cartoon schematic of artificial decidualization model. Image on right demonstrates difference in size in oil-treated versus control uterine horns. (D-G) Immunofluorescence images of *Nestin*-CreER; *Rosa*-Tomato uteri 4 days, 8 days, and 10 days post-surgery for artificial decidualization. Arrows indicate Tomato+ cells in the epithelium. (H) Quantification of percent Tomato-positive epithelial cells in oil-treated versus control uterine horns 10 days post-surgery. Thin scale bar: 200 μm; thick scale bar: 100 μm. \*\*\**P*<0.001.

If pregnancy does not occur, the decidualized endometrium is shed during menstruation. Unlike in humans, mouse endometrium undergoes decidualization in implantation sites during pregnancy^4^. To mimic human endometrial regeneration after menstruation, artificial decidualization in mice is induced by a single injection of sesame oil in the pseudo-pregnant uterine lumen^38^. In this artificial decidualization model, decidualized stromal cells together with original luminal epithelial cells are shed from the uterus, and a new epithelial lining forms under the detached decidual tissues (Fig. 3C). To investigate the role of *Nestin*+ perivascular cells in regenerating the new epithelial lining, we collected uterine tissues 4, 6, 8 and 10 days after oil injection, in which the decidual reaction peaks 4 days after oil injection and the original luminal epithelium has degenerated. Tomato+ cells were recruited adjacent to the original lumen, with some of them expressing CDH1 (Fig. 3D), but these cells will eventually be shed together with all decidual cells. Short sections of new epithelial lining began to emerge between the decidualized and intact stromal regions on the periphery of the section at four days after oil injection (Fig. 3D, right panel). Six days after oil injection, the border between the decidualized and intact stromal regions became more prominent, and an empty space was formed due to the gradual detachment of the decidualized tissues (Fig. 3E). The new epithelial lining covering the intact stromal tissues became more polarized as evident by the cuboidal shapes of epithelial cells. Notably, Tomato+ cells were observed in the newly regenerated epithelium, while more Tomato-labeled cells were in close proximity subjacent to the epithelium (Fig. 3E). Eight days post-injection, the new epithelial lining was completely formed, with decidualized tissues shed into the lumen waiting to be expelled. Clones of Tomato+ epithelial cells were observed in multiple loci within the new epithelial lining (Fig. 3F). By 10 days after oil injection, all decidualized tissues were removed and the endometrial tissue was almost fully restored (Fig. 3G). Quantification revealed that about 12% of epithelial cells in the oil-injected uterine horn was Tomato-positive, while a significantly lower percentage (∼1%) was Tomato-positive in the control un-injected uterine horn (Fig. 3H). Taken together, these results suggest that the contribution of *Nestin*+ perivascular cells to epithelial regeneration is increased during circumstances of significant epithelial loss.

### Nestin-derived cells undergo mesenchymal-to-epithelial transition *in vitro*

Our *in vivo* lineage-tracing analyses suggested that uterine Nestin+ perivascular cells undergo mesenchymal-to-epithelial transition (MET). To further investigate the ability of Nestin-derived perivascular cells to undergo MET and to directly observe their differentiation into epithelial cells, we employed an *in vitro* endometrial organoid (EMO) culture system, which closely approximates the physiology of the uterus^39^. For organoid assays, we FACS-purified Tomato+ cells from *Nestin*-CreER; *Rosa*-Tomato endometrium 2 days after exposure to 4-OHT at P60. Whole endometrial cells from the same mouse model served as controls. After 6 days in culture following an established protocol^39^, both whole uterine cells and FACS-purified Tomato+ cells self-organized into organoid-like structures (Fig. S3A). Very few Tomato+ cells were observed in control organoids, perhaps due to epithelial cells being more adept or efficient at forming EMOs compared to other cells (Fig. S3A). Despite being able to readily form EMOs, the number of organoids from purified Tomato+ cells was significantly lower than controls (Fig. S3B), and Tomato+ organoids were smaller in size (Fig. S3C). The cells in all organoids from both groups expressed ERα and the epithelial marker CDH1 (Fig. S3D), suggesting *Nestin*+ perivascular cells can differentiate into epithelial cells and are capable of assembling EMOs.

### Estrogen triggers the differentiation of *Nestin*+ perivascular cells into epithelial cells

The remodeling of endometrium is orchestrated by fluctuating levels of estrogen and progesterone^3^; therefore, we examined the expression of estrogen and progesterone receptors in uterine perivascular cells. Two months post-exposure to 4-OHT at P60, Tomato-labeled cells expressed ERα and PGR in *Nestin*-CreER; *Rosa*-Tomato endometrium (Fig. 4A and C). In an artificial decidualization model, Tomato+ cells in the stroma and the newly regenerated epithelium also expressed ERα and PGR (Fig. 4B and D). These findings suggest that uterine *Nestin*+ perivascular cells can respond to estrogen and progesterone signals.

**Figure 4.**
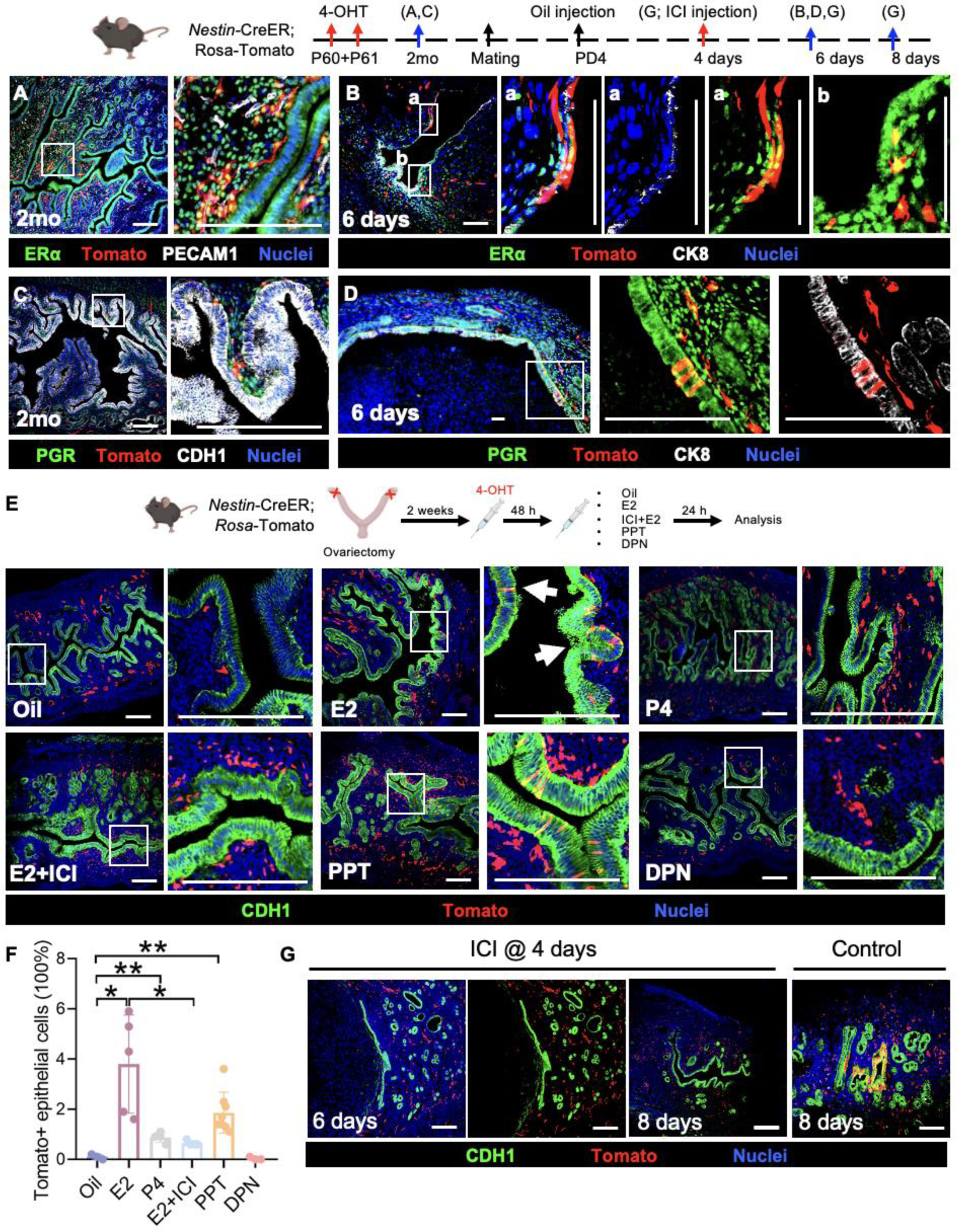
Estrogen triggers the differentiation of *Nestin*+ perivascular cells into epithelial cells via ERα. (A-D). Immunofluorescence images of *Nestin*-CreER; *Rosa*-Tomato uteri at P62 and 6 days post-surgery for artificial decidualization following administration of 4-OHT at P60 and P61. (E) Immunofluorescence images of *Nestin*-CreER; *Rosa*-Tomato uteri after ovariectomy and various hormonal treatments. (F) Quantification of percent Tomato-positive epithelial cells after various hormonal treatments. (G) Immunofluorescence images of *Nestin*-CreER; *Rosa*-Tomato uteri 6 and 8 days post-surgery for artificial decidualization. Scale bar: 100 μm. \**P*<0.05, \*\**P*<0.01.

To determine the effect of estrogen and progesterone on perivascular cell fate, ovariectomized *Nestin*-CreER; *Rosa*-Tomato females were allowed to recover for 2 weeks to eliminate any remaining ovarian hormones and were then administered 4-OHT to label *Nestin*+ perivascular cells. After 48 hours, a single injection of estradiol (E2, 100 ng/mouse) or progesterone (P4, 1 mg/mouse) was administered to induce a specific hormonal response as previously described^40,41^. After 24 hours, Tomato-labeled cells were observed within the epithelium of uteri exposed to E2, while very few Tomato+ cells were observed following P4 injection (Fig. 4E and F). To determine if this differentiation was estrogen-receptor-dependent, an estrogen receptor antagonist, ICI 182,780 (0.5 mg/mouse), was administered 1 hour before E2 injection. Tomato-labeled cells were not detected in the epithelium after estrogen receptor blockade (Fig. 4E and F). Next, PPT (an ERα agonist, 250 ng/mouse) or DPN (an ERβ agonist, 100 ng/mouse) were given to determine the estrogen receptor mediating the effect of E2. We observed more Tomato+ epithelial cells in uteri exposed to PPT than those in the DPN-treated group, the latter of which was comparable to control, non-treated uteri (Fig. 4E and F). These results demonstrate that estrogen promotes the differentiation of Tomato+ perivascular progenitors into epithelial cells via ERα.

To further test whether ERα is required for the differentiation of perivascular cells during endometrial remodeling, we injected a single dose of the estrogen receptor antagonist ICI 182,780 at 4 days post-oil injection in an artificial decidualization model. Two days after ICI 182,780 injection, no Tomato-labeled cells were observed in the endometrial epithelium (Fig. 4G). Taken together, these results indicate that the differentiation of perivascular cells occurs in an estrogen/ERα-dependent manner.

### Estrogen suppresses Notch activity in perivascular cells via ERα

How estrogen initiates the differentiation of progenitors during endometrial regeneration is poorly understood. A potential player in this process is Notch signaling, which is indispensable for maintaining *Nestin*+ perivascular cells in developing gonads^35,36^. Using *CBF:*H2B-Venus^42^ reporter mice, which express Venus upon Notch activation, we found that Notch signaling was active in perivascular cells expressing ERα in P60 endometrium (Fig. 5A). However, few perivascular cells underwent active Notch signaling without expressing nuclear ERα (Fig. 5A). To determine the regulation of Notch activity by estrogen, ovariectomized *CBF*:H2B-Venus reporter females were exposed to estradiol, ICI (estrogen receptor antagonist) or estrogen receptor activators (Fig. 5B). Estrogen significantly decreased uterine Notch activity, but this downregulation of Notch activity was blocked by ICI, suggesting that estrogen suppressed Notch signaling via ERs. Next, we found that perivascular Notch activity was inhibited by PPT injection but not DPN, indicating that estrogen mediated this suppression of Notch via ERα (Fig. 5C). qRT-PCR analyses showed that mRNA levels of the Notch target genes *Hes1*, *Hes5*, and *Hey2* were all significantly down-regulated by both estrogen and PPT, but ICI blocked this downregulation (Fig. 5D). We found that both *Notch2* and *Notch3* were downregulated by estrogen and PPT, while *Notch1* and *Notch4* levels remained unaffected (Fig. 5E). The mRNA levels of *Dll4* also exhibited a similar trend, while *Jagged2* was unaffected and *Dll1* was not downregulated by PPT (Fig. 5F). *Notch2* was enriched in various cell types in D4 uterus based on published single-cell RNA-seq data^43,44^, while *Notch3* was specifically enriched in pericytes and *Dll4* was enriched in endothelial cells (Fig. S4A-C). On day 8 of pregnancy, both *Notch3* and *Dll4* were expressed in the same cell types as on day 4 (Fig. S4D and E). Furthermore, to investigate whether Notch activity in perivascular cells is maintained by Notch3, we examined Notch3 expression in *CBF*:H2B-Venus mice and found that perivascular Venus+ cells expressed Notch3 (Fig. 5G). We observed that DLL4 was expressed in endothelial cells, which were adjacent to Venus+ cells (Fig. 5G). These data suggest that perivascular cells undergo active Notch signaling, potentially through DLL4-Notch3 interactions, which can be suppressed by ERα-mediated E2 signaling.

**Figure 5.**
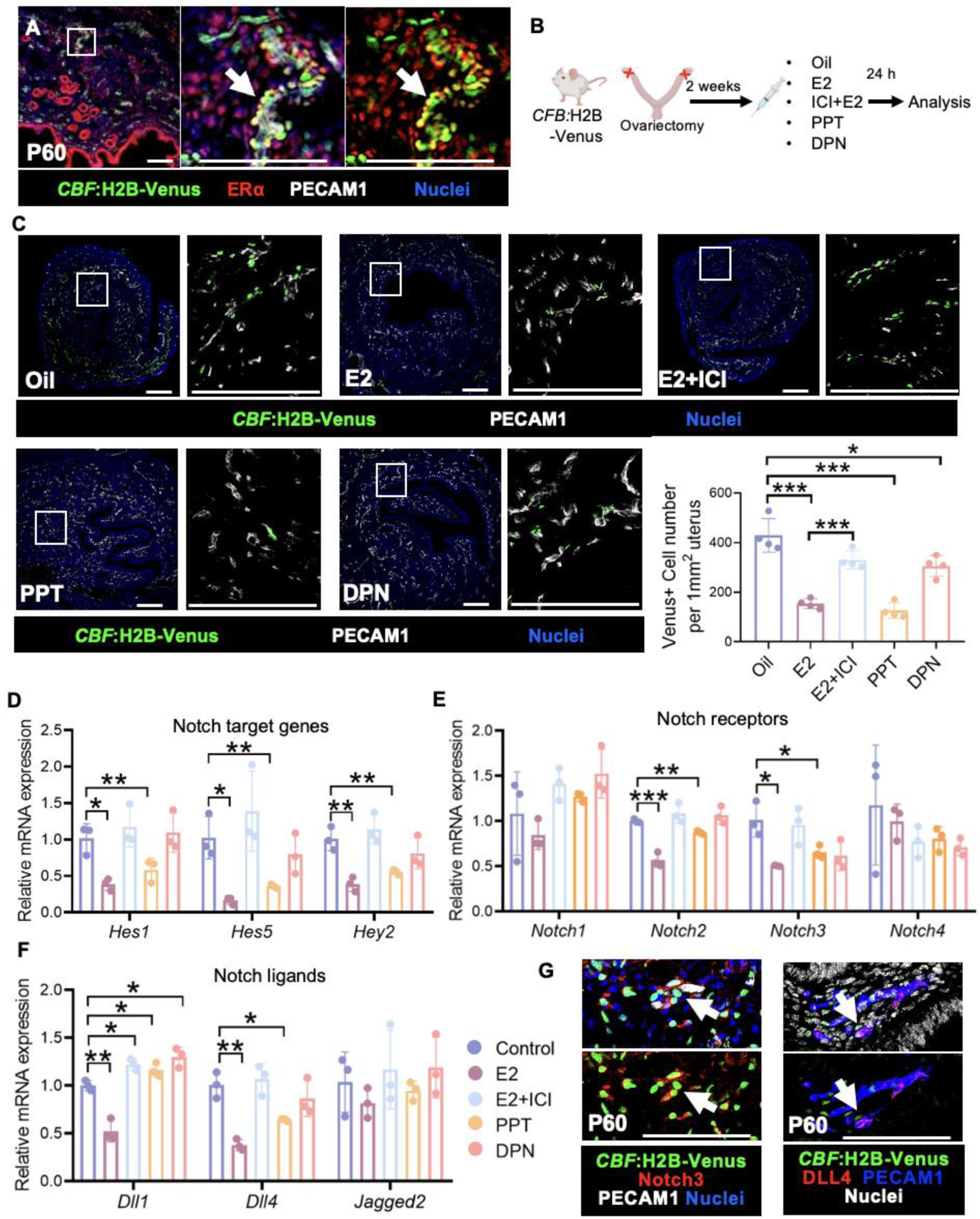
Estrogen suppresses Notch activity in the uterus. (A) Immunofluorescence images of P60 *CBF*:H2B-Venus uterus. Arrow indicates Venus-expressing cells adjacent to the vasculature that express ERα. (B) Scheme of ovariectomized mouse model and treatment. (C) Immunofluorescence images of *CBF*:H2B-Venus uteri post ovariectomy after various hormonal treatments. (D-F) qRT-PCR analyses showing mRNA levels of Notch target genes (D), Notch receptors (E), and Notch ligands (F). (G) Immunofluorescence images of P60 *CBF*:H2B-Venus uteri. White arrows point to perivascular Venus-expressing cells that express Notch3, and yellow arrows point to Venus-expressing cells adjacent to DLL4+ vasculature. Scale: 100 μm. \**P*<0.05, \*\**P*<0.01, \*\*\**P*<0.001.

### Notch signaling is required for maintaining homeostatic quiescence of *Nestin*+ perivascular cells

To determine if perivascular *Nestin*+ cells undergo active Notch signaling, we examined Nestin expression in P60 and D8 endometria of *CBF:*H2B-Venus reporter mice and observed that Venus-expressing perivascular cells also expressed Nestin (Fig. 6A and B). Furthermore, perivascular Tomato-labeled cells in P62 *Nestin*-CreER; *Rosa*-Tomato uteri expressed Notch3 and were adjacent to cells expressing DLL4, which was specifically expressed in endothelial cells (Fig. 6C and D). Using *Nestin*-CreER; *Rosa*-Tomato; *CBF:*H2B-Venus mice, we observed that perivascular Tomato-labeled cells were actively involved in Notch signaling in non-pregnant uteri (Fig. S5A). During epithelial regeneration, perivascular Tomato+ cells underwent active Notch signaling (Fig. S5B and C-c) and continued to exhibit active Notch signaling in the regenerating epithelium (Fig. S5C-b). However, active Notch signaling was not observed in these cells once the epithelium had fully regenerated (Fig. S5C-c). These data suggest that Notch activity is involved in maintaining the undifferentiated state of perivascular cells.

**Figure 6.**
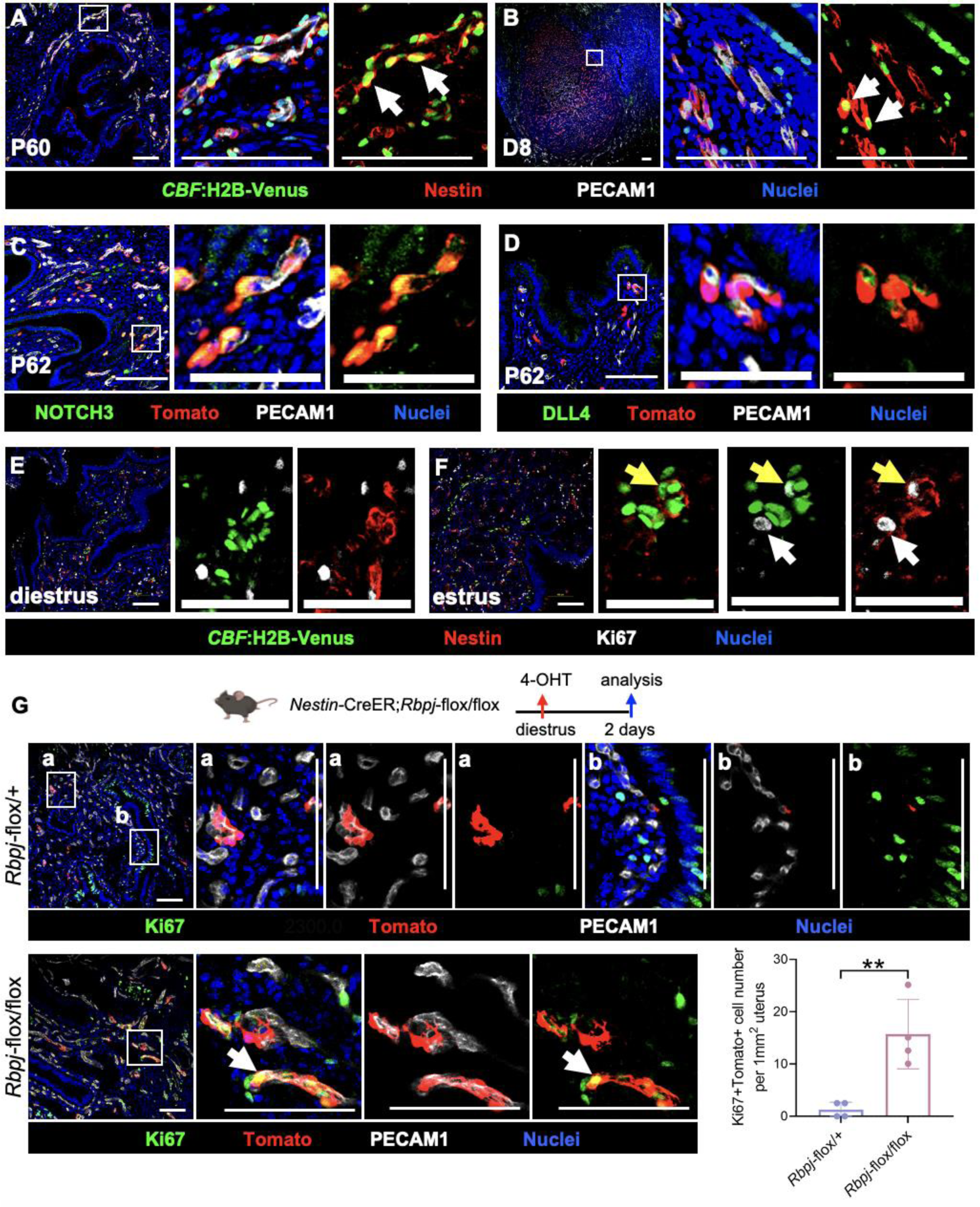
Notch activity maintains quiescence of *Nestin*+ perivascular cells. (A-B) Immunofluorescence images of P62 and D8 *CBF*:H2B-Venus uteri. Arrows point to cells co-expressing Venus and Nestin. (C-D) Immunofluorescence images of P62 *Nestin*-CreER; *Rosa*-Tomato uteri. (E-F) Immunofluorescence images of *CBF*:H2B-Venus uteri at diestrus and estrus phases. White arrows indicate Nestin-expressing cells without Venus expression that express Ki67, and yellow arrows point to Nestin-expressing cells with low Venus expression that express Ki67. (G) Immunofluorescence images of *Nestin*-CreER; *Rosa*-Tomato; *Rpbj*-flox/+ heterozygous and *Nestin*-CreER; *Rosa*-Tomato; *Rpbj*-flox/flox uteri 2 days after exposure to 4-OHT at the diestrus phase. Arrow indicates cell that co-expresses Ki67 and Tomato. Thin scale bar: 200 μm; thick scale bar: 100 μm. Graph shows number of Ki67+Tomato+ cells per unit area of uterus. \*\**P*<0.01.

Maintaining cell cycle quiescence of adult stem cells is crucial for life-long tissue homeostasis and regenerative capacity^45^. In *Nestin*-CreER; *Rosa*-Tomato uteri previously exposed to 4-OHT (2 months prior), most Tomato+ cells were Ki67-negative during the diestrus phase, while some Tomato+ cells were Ki67-positive during the estrus phase (Fig. S5D). Quantification analyses indicated that the number of Ki67+Tomato+ cells was significantly increased in estrus-phase uteri compared to diestrus-phase uteri (Fig. S5E). We observed that almost all Nestin+ perivascular cells undergoing active Notch signaling did not express Ki67 in the diestrus phase of *CBF*:H2B-Venus reporter uteri, indicating a state of quiescence (Fig. 6E). However, during the estrus phase, a subset of Nestin+ perivascular cells with low or no Notch activity expressed Ki67, suggesting a transition toward an active cell cycle state (Fig. 6F). Taken together, these results indicate that *Nestin*+ perivascular cells exhibiting active Notch activity remained quiescent in non-pregnant uterus, while those that displayed reduced or no Notch activity during the estrus phase entered an active cell cycle state.

To address whether Notch signaling is required in perivascular cells to maintain their quiescence, we performed a cell-type–specific conditional deletion of the Notch transcriptional regulator *Rbpj* (also known as *Cbf1*) using a *Nestin*-CreER; *Rosa*-Tomato; *Rpbj*-flox/flox mouse model. Adult control *Nestin*-CreER; *Rosa*-Tomato; *Rpbj*-flox/+ heterozygous and *Nestin*-CreER; *Rosa*-Tomato; *Rpbj*-flox/flox mice were exposed to 4-OHT during the diestrus phase and uteri were collected after 48 hours. We detected RBPJ protein in significantly fewer Tomato+ cells in *Nestin*-CreER; *Rosa*-Tomato; *Rpbj*-flox/flox uteri as compared to heterozygous controls (Fig. S5F and G), demonstrating the effectiveness of the deletion. Tomato+ cells in control uteri were overwhelmingly perivascular, undifferentiated, and Ki67-negative (Fig. 6G). In *Rbpj* conditional knockout uteri, there was an increase in the number of perivascular Tomato+ cells that were Ki67-positive (Fig. 6G). These results suggest that *Notch* signaling is required to maintain quiescence of *Nestin*+ perivascular cells.

### Estrogen induces cell cycle activity in *Nestin*+ perivascular cells via inhibiting Notch activity

Quiescent stem/progenitor cells enter the cell cycle when activated in response to extrinsic signals, whereby the fate of multipotent progenitors is usually determined in the G1 phase of the cell cycle^47^, potentially leading to differentiation, senescence, or re-entry into quiescence^47^. To investigate whether estrogen induces cell cycle activation in *Nestin*+ perivascular cells, we examined Ki67, which is expressed during all active cell-cycle phases but not in the resting phase G0^48^, in ovariectomized *Nestin*-CreER; *Rosa*-Tomato females. Tomato+ cells were Ki67-negative in controls, whereas the number of Tomato+ Ki67+ cells was significantly increased after estrogen exposure for 24 hours (Fig. 7A, B). To determine if estrogen activated these perivascular cells by suppressing Notch activity, we examined Ki67 in ovariectomized *CBF*:H2B-Venus reporter mice after 24 hours of estradiol exposure. In controls, we observed most Nestin+ cells were undergoing active Notch signaling and rarely expressed Ki67. In contrast, estrogen treatment decreased Venus expression and induced Nestin+ cells to express Ki67 (Fig. 7C).

**Figure 7.**
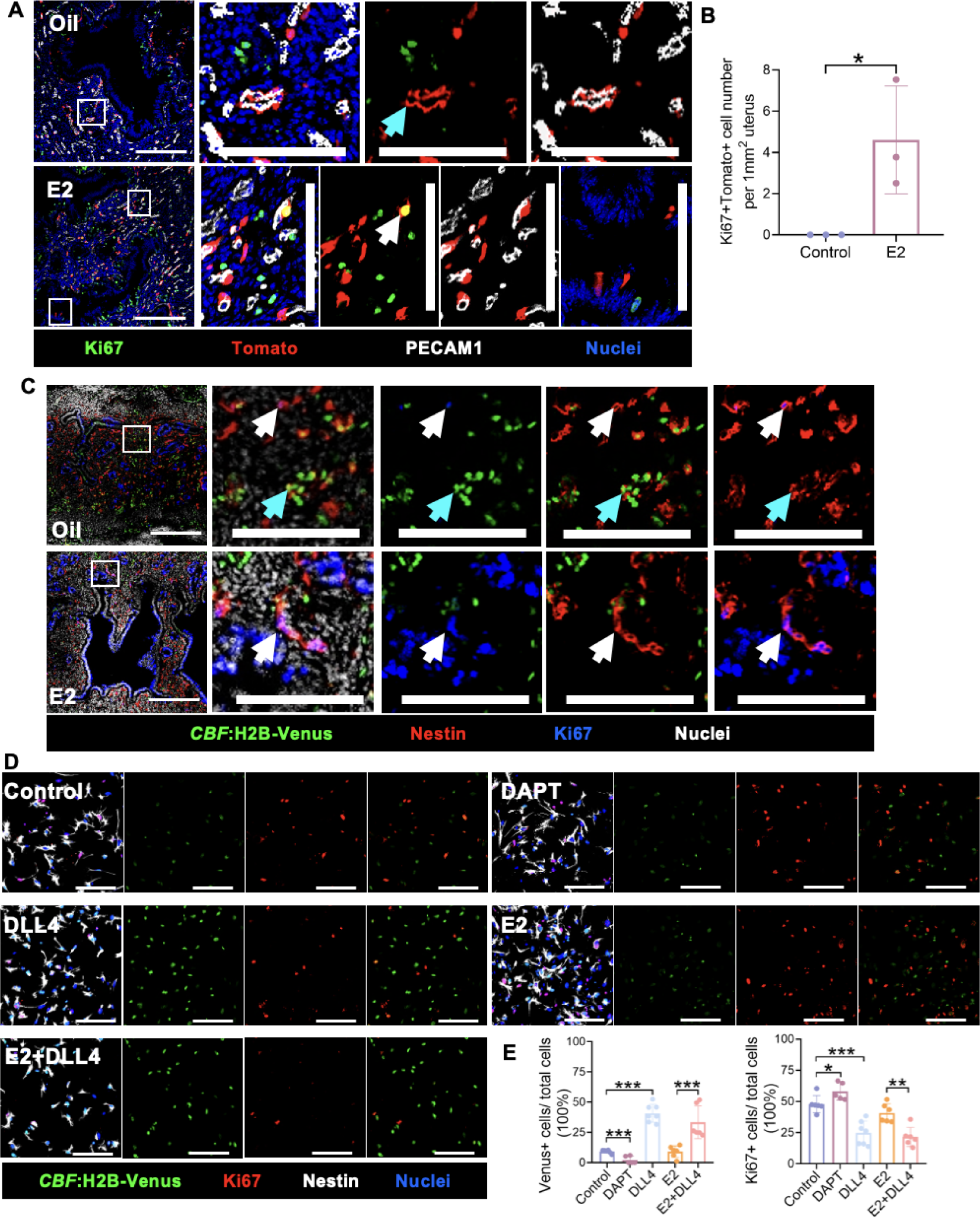
Estrogen activates quiescent *Nestin*+ perivascular cells via inhibiting Notch activity. (A) Immunofluorescence images of ovariectomized *Nestin*-CreER; *Rosa*-Tomato uteri 24 hours after exposure to oil control or estrogen. White arrow indicates Tomato+ cell that expresses Ki67, while cyan arrow indicates Ki67-negative Tomato+ cell. (B) Quantification of Ki67+Tomato+ cell number per unit area. (C) Immunofluorescence images of ovariectomized *CBF*:H2B-Venus uteri 24 hours after exposure to oil control or estrogen. White arrow indicates Ki67+Nestin+ cell without Venus expression, while cyan arrow indicates Ki67+Nestin+ cell that co-expresses Venus. (D) Immunofluorescence images of Venus+ cells from P60 *CBF*:H2B-Venus endometrium after *in vitro* culture for 4 days in various treatment conditions. Thin scale bar: 200 μm; thick scale bar: 100 μm. (E) Quantification of percent Venus+ (left) and Ki67+ (right) cells in *in vitro* culture assays. \**P*<0.05, \*\**P*<0.01, \*\*\**P*<0.001.

To further examine if estrogen activated Nestin+ cells by inhibiting Notch signaling, stromal Venus+ cells were disassociated from *CBF*:H2B-Venus uteri and treated with E2 and/or DLL4. After 4 days, most cells lost Venus expression but almost all expressed Nestin (Fig. 7D), suggesting that these cells were *Nestin*+ cells and Notch signaling was inactivated once they left the perivascular niche. In culture on DLL4-coated slides, the expression of Venus was significantly increased, suggesting that Notch signaling was successfully activated and/or maintained. Moreover, we observed that the number of Ki67+ cells was significantly decreased in DLL4-treated cells compared with controls, indicating that activation of Notch signaling maintained the quiescence of these cells *in vitro* (Fig. 7D and E). Taken together, the results from *in vivo* and *in vitro* assays suggest that estrogen activates quiescent *Nestin*+ cells to enter the cell cycle by inhibiting Notch signaling.

## Discussion

The dynamic nature and regenerative capacity of the endometrium are critical for the establishment and maintenance of successful pregnancies^1,2^. A remarkable regenerative capacity indicates the presence of stem/progenitor cells in the endometrium, but their precise identification has been elusive, particularly within the stroma. Perivascular cells in the endometrium are one cell type speculated to be mesenchymal stem cells. They share phenotypic characteristics with mesenchymal stem cells, indicating their potential in endometrial regeneration and repair^9,17^. Our previous research showed that *Nestin*+ perivascular cells in the gonads serve as progenitors: in the testis, they become Leydig cells and other mesenchymal-derived cell types, and in the ovary, they differentiate into granulosa cells as well as interstitial cell types^35,36^. Here, using a *Nestin*-CreER; *Rosa*-Tomato mouse model, we show that *Nestin*+ perivascular cells contribute to endometrial epithelial regeneration during estrous cycles and pregnancy (Fig. 8). While these cells are usually in a quiescent state within a perivascular niche maintained by Notch activity, estrogen induces them to exit quiescence by suppressing Notch activity. Estrogen, acting via ERα expressed on these perivascular cells, mediates their transformation into epithelial cells. Endometrial epithelial regeneration has been presumably attributed to *in situ* proliferation of resident or residual epithelial cells. Our study identifies *Nestin*+ perivascular cells as epithelial progenitor cells participating in endometrial epithelial regeneration. This discovery significantly advances our understanding of uterine regeneration and opens up a new avenue of research in maintaining reproductive health for women of childbearing age.

**Figure 8.**
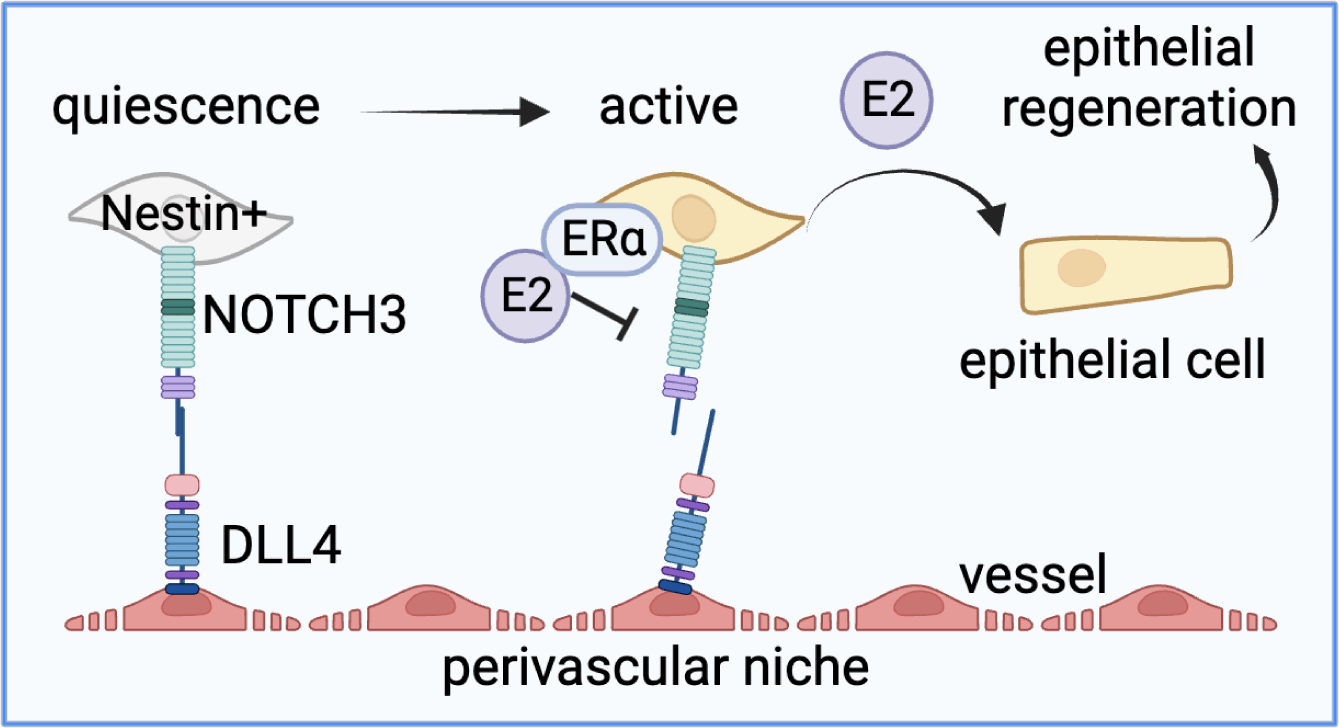
A perivascular niche regulates epithelial progenitors in the uterine stroma. Cartoon model depicts how estrogen directs the quiescence and activation of *Nestin*+ perivascular cells in the uterus through modulation of vascular-mesenchymal Notch signaling interactions.

Adult stem cells have the remarkable potential to develop into different cell types. Compared to embryonic stem cells, they have a more limited ability to differentiate into various cell types but play a vital role in maintaining and repairing the tissue in which they reside^49^. Adult stem cells can be categorized based on their origin and differentiation propensity, including hematopoietic stem cells, mesenchymal stem cells, neural stem cells, and epithelial stem cells^50-53^. Endometrial stem cells endow regenerative properties that enable the restoration of the endometrial epithelial layer after menstruation or during the postpartum period^54^. In particular, postpartum and post-menstrual endometrial repair involve rapid re-epithelialization. Classical studies suggested that newly regenerated epithelia are mainly derived from proliferation of the remaining epithelia close to denuded stromal cells^55^. SSEA-1+ epithelial stem cells serve as a resident pool of progenitors for epithelial regeneration in uterus^6,16,37,54,56^. They have the potential to proliferate and restore epithelial integrity in case of cell death nearby^8^. Additionally, recent observations of stromal cells that expressed both cytokeratin and vimentin close to the denuded surface suggests that mesenchymal-to-epithelial cell trans-differentiation (MET) could be another source of newly restored epithelial cells^9,10^. Using *Amhr2*-Cre transgenic mice to lineage trace stromal cells and their progeny, previous studies showed that *Amhr2*+ stromal cells contribute to endometrial re-epithelialization after menses and delivery^14,15^. To confirm the involvement of MET in endometrial repair, researchers labeled all mesenchymal cells using *Pdgfrb-*rtTA or stromal fibroblasts using *Pdgfra-*CreER in adulthood, but results were seemingly contradictory. The study using *Pdgfrb-*rtTA to trace mesenchymal cells did not find any incorporation of labeled cells in new epithelia^57^, whereas stromal cells labeled by *Pdgfra-*CreER contributed to endometrial re-epithelization using a mouse artificial menstrual model^16^. These findings indicated the possible presence of adult mesenchymal stem/progenitor cells as a source for endometrial epithelial regeneration. But the precise cell type remains unidentified due to the heterogeneity of endometrial mesenchymal cells, including stromal fibroblasts, vascular, and perivascular cells^58,59^.

Uterine perivascular cells express mesenchymal-stem-cell-specific genes^18^. However, whether they exhibit functional stem cell properties remains unclear. Recently, a recent single-cell study reported that perivascular cells rapidly expand during endometrial regeneration using a mouse menstruation model, but they disappear once endometrial repair is complete. Therefore, perivascular cells were identified as a repair-specific cell type that contributes to endometrial repair. However, an *Ng2*-CreER mouse model suggested that *Ng2*+ perivascular cells did not contribute to epithelial repair^16^. Considering that there are at least two subpopulations of perivascular cells in non-pregnant uterus and one newly regenerated subpopulation during endometrial repair^16^, the *Ng2*-CreER mouse model might not target all subtypes of endometrial perivascular cells. In the current study, we found that Nestin protein was specifically expressed in perivascular cells in the uterus. During the estrous cycle, these perivascular cells not only maintained their Nestin expression but also contributed to epithelial expansion, with a distinct subpopulation emerging adjacent to the uterine epithelium, showing selective Nestin expression (and not NG2). These findings suggest that there may be unique sets of pericytes with varying functions within the uterus. Interestingly, the emergence of a distinct subpopulation of perivascular cells adjacent to the uterine epithelium further underscores the complexity and heterogeneity of the endometrial cellular landscape. The dynamic behavior of Nestin and NG2, especially during the early stages of pregnancy, reveals their potentially significant and unique roles in dynamic uterine changes during pregnancy. The consistent expression of Nestin during early pregnancy and its spatial distribution pattern in the uterus suggest that these cells have a long-lasting presence and do not undergo drastic changes during pregnancy, unlike NG2+ cells. Using pregnancy and mouse artificial decidualization models, we examined the role of *Nestin*+ perivascular cells in endometrial epithelial regeneration. We found that about 10% of the endometrial epithelium is derived from these cells after endometrial repair. However, due to the low Tomato labeling efficiency in *Nestin*-CreER; *Rosa*-Tomato uteri, this contribution is likely significantly underestimated. We attempted to enhance this efficiency by increasing either the 4-OHT dose or the number of exposure days to 4-OHT. However, these regimens led to undesirable side effects: peritoneal adhesions, infertility, and failure to induce decidualization. Therefore, it is technically challenging to assess fully the contribution of perivascular cells during uterine regeneration using this particular mouse model.

Adult stem cells reside in specialized microenvironments known as niches, which are critical for maintaining stem cell properties and function^60^. Among these, the perivascular niche is particularly significant and unique, characterized by interactions between endothelial cells and perivascular cells that regulate stem cell behavior^61^. Departure from this niche often triggers differentiation in stem cells. Our previous studies found that disruption of vascular endothelial cells induced the differentiation of perivascular cells in fetal gonads, indicating that the perivascular niche plays a major role in maintaining mesenchymal stem cell identity within reproductive tissues^36,62^. A pathway that is likely involved is Notch, which is a key pathway for maintaining the quiescence and homeostasis of mesenchymal stem cells^25,36,62^. Endothelial cells provide Notch ligands that initiate the Notch signaling cascade, while perivascular cells express Notch receptors that receive and transduce these signals. Disruption of Notch-ligand interactions can alter stem cell fate and has profound implications for tissue homeostasis and repair^63^. In fetal gonads, endothelial cells maintain undifferentiated progenitors in a perivascular niche via Notch signaling, and blocking Notch signaling prompts perivascular cells to differentiate into Leydig cells or granulosa cells in testes and ovaries, respectively^36,62^. In the uterus, Nestin+ perivascular cells strongly expressed Notch3, while DLL4, a Notch ligand, was predominately expressed in endothelial cells. Using *CBF*:H2B-Venus Notch reporter mice, we found that most Nestin+ cells undergoing active Notch signaling were in the resting/quiescent phase of the cell cycle, G0 (Ki67-negative), during the diestrus phase. In contrast, Nestin+ cells with low or no Notch activity entered the cell cycle (Ki67+) during the estrus phase. However, in perivascular-specific conditional knockout mice for the Notch transcription factor *Rbpj*, *Nestin*+ perivascular cells did not undergo differentiation or leave their perivascular niche, but instead transitioned from a quiescent to an active state in the uterus. In human endometrial mesenchymal stromal/stem-like cells (CD140b+/CD146+), activation of Notch signaling via JAG1-NOTCH1 interaction is required for maintaining their quiescent state *in vitro*^25^. Adult stem cells stay in a quiescent state to prevent the loss of self-renewal capacity and to avoid the rapid exhaustion of adult stem cells^45^. Notch signaling is also essential for the sustained preservation of adult neural stem cells. When Notch signaling is inactive, neural stem cells that divide slowly start to transform into transit-amplifying cells and eventually into neurons. This results in a temporary boost in the generation of new neurons. However, over time, this leads to the exhaustion of the neural stem cell population, resulting in a reduction of neuron formation^64^. Inhibition of Notch signaling via uterine cavity injection of DAPT, a γ-secretase inhibitor, delays endometrial repair following menstrual-like breakdown in mice^25^. This suggests that modulating adult stem cell activity via Notch is critical for endometrial repair, and any disruption of this balance compromises the repair process.

Most uterine *Nestin*+ perivascular cells were maintained in a quiescent state during the diestrus phase, while some of them entered the active cell cycle during the estrus phase, suggesting that estrogen might activate quiescent perivascular cells. Using an ovariectomized *Nestin*-CreER; *Rosa*-Tomato mouse model, we found that estrogen activated the cell cycle of quiescent Nestin+ perivascular cells. In breast cancer, estrogen controls a variety of genes, such as protooncogenes and growth factors, that induce cell cycle activity. Additionally, estrogen directly influences proteins that govern the cell cycle and also enhance the activity of proteins that promote cell cycle arrest^65^. Our analyses using ovariectomized *CBF*:H2B-Venus reporter mice show that estrogen could reduce uterine Notch activity. Nestin+ cells usually remained inactive within a perivascular niche through Notch signaling interactions. In our ovariectomized *CBF*:H2B-Venus reporter mouse model, estrogen exposure led to a decrease in Notch activity in Nestin+ cells. In contrast, some Nestin+ cells with low or no Notch activity expressed Ki67, suggesting they were no longer quiescent and entered an active state. These findings indicate that estrogen drives the activation of Nestin+ cells by inhibiting Notch activity in the uterus.

Although Notch signaling is crucial for maintaining stem cells in a quiescent state, cellular differentiation is dependent on other signaling mechanisms. Endometrial regeneration is orchestrated by circulating estrogen and progesterone, in which estrogen is a critical factor for this regeneration^3,66^. Estrogen stimulates the expansion of endometrial epithelium and stroma in ovariectomized mice via ERα^41,67,68^. Conditional deletion of *Esr1* (encoding ERα) in stromal cells using an *Amhr2*-Cre; *Esr1*-floxed mouse model showed defects in estrogen-induced epithelial expansion^69^. Using *Nestin*-CreER; *Rosa*-Tomato mice, we observed that uterine *Nestin*+ perivascular cells expressed ERα in non-pregnant and pregnant endometrium. This expression indicates that these cells have the potential to respond to estrogen signaling. In ovariectomized mice, both estrogen and an ERα agonist triggered the differentiation of *Nestin*+ perivascular cells into epithelial cells, while an estrogen receptor inhibitor hindered this differentiation. Furthermore, we did not observe any Tomato-labeled *Nestin*+ cells in newly regenerated epithelium in *Nestin*-CreER; *Rosa*-Tomato endometrium treated with estrogen receptor inhibitor during epithelial regeneration in a mouse menstruation model. Taken together, these results indicate that estrogen induces the differentiation of *Nestin*+ perivascular cells into epithelial cells in an ERα-dependent manner. Interestingly, a previous study showed that estrogen is not required for uterine regeneration^70^. It is possible that the participation of Nestin+ cells in endometrial regeneration is limited, and, instead, proliferation of remnant epithelial cells is the main source of cells for regeneration. Estrogen regulates the differentiation process of neural stem cells, dendritic cells, and early osteoblasts^71-73^. In the endometrium, estrogen directly induces the differentiation of other cells, including uterine natural killer cells (uNK) in the decidual environment. Estrogen binds to its receptors to activate expression of immunomodulatory or angiogenic proteins in uNK cells, or to increase expression of alectin-1, an immunosuppressant, in the decidual NK population^74,75^. Further investigation is required to understand how estrogen triggers the differentiation of perivascular cells into epithelial cells.

In conclusion, our study uncovers a previously unappreciated role for perivascular cells as adult stem/progenitor cells that participate in endometrial regeneration, whereby these cells differentiate into epithelial cells in response to estrogen via ERα. Our study provides valuable insights into the cellular and molecular mechanisms that underpin the regeneration of the endometrial lining after each menstrual cycle and during postpartum recovery. The interplay between estrogen and Notch activity emerges as a critical regulatory mechanism in directing the quiescence and activation of these cells. Understanding the mechanisms of quiescence maintenance and fate decision in perivascular stem cells provides insight into endometrial cell fate decisions, informing new potential therapeutic strategies for uterine disease.

## Methods

### Mice

Cre-responsive *Rosa*-Tomato [B6.Cg-*Gt(ROSA)26Sor^tm14(CAG-tdTomato)Hze^*; JAX stock #007914] and *CBF*:H2B-Venus [Tg(Cp-HIST1H2BB/Venus)47Hadj; JAX stock #020942] mice were obtained from the Jackson Laboratory. *Nestin*-CreER mice [Tg(Nes-cre/ERT2,-ALPP)1Sbk]^34,35^ were obtained from Masato Nakafuku, Division of Developmental Biology, Cincinnati Children’s Hospital Medical Center. *Rbpj*-floxed mice (*Rbpj*^tm1Hon^)^76^ were provided by Joo-Seop Park, Division of Pediatric Urology, Cincinnati Children’s Hospital Medical Center. *Nestin*-CreER; *Rosa*-Tomato mice in diestrus cycle were injected intraperitoneally with 4-hydroxytamoxifen (4-OHT, Sigma-Aldrich, #H6278, St. Louis, MO) to induce CreER recombinase activity. For two consecutive days, each mouse received a dose of 2 mg of 4-OHT (injected intra-peritoneally), which was initially dissolved in ethanol and then mixed with corn oil. *Nestin*-CreER-negative; *Rosa*-Tomato mice, which did not express the CreER recombinase, were used as a control group to account for any background fluorescence and to assess any potential leakiness of Tomato reporter mice. Mice were housed in accordance with National Institutes of Health guidelines, and experimental protocols were approved by the Institutional Animal Care and Use Committee (IACUC) of Cincinnati Children’s Hospital Medical Center (protocol #IACUC2021-0016).

### Identification of estrous cycle

The estrous cycle is divided into four phases, namely proestrus, estrus, metestrus and diestrus, and lasts for 4 to 5 days. Staging of the estrous cycle was performed as described^77,78^. Briefly, vaginal swabs were collected from mice. Cells on the swab were transferred to a glass slide and air-dried. After air-drying, slides were stained with 4% Giemsa solution for 10 min at room temperature and rinsed with water. The estrous cycle stage was manually determined based on the percentages of leukocytes, cornified epithelial cells, and nucleated epithelial cells.

### Artificial decidualization

Wild-type and transgenic females were mated with vasectomized wild-type males to induce pseudo-pregnancy. One uterine horn was intra-luminally injected with 50 μl sesame oil under anesthesia at pseudo-pregnancy day 4 (PD4) as previously described^38^; the other uterine horn was left untreated as a control. Uteri were collected at day 8, day 10, and day 12 of pseudo-pregnancy (critical time points in the regeneration process) for subsequent downstream analyses.

### Hormone treatment

Eight-week-old mice were ovariectomized and allowed to recover for two weeks to ensure clearance of any residual ovarian hormones. A single injection of estradiol-17β (E2, 100 ng/mouse, E1024, Sigma Aldrich, St. Louis, MO) was injected subcutaneously to induce a specific hormonal response as previously described^40^. Estrogen receptor antagonist ICI 182,780 (0.5 mg/mouse, Sigma Aldrich, #V900926, St. Louis, MO) was administered 1 hour before E2 injection to block all estrogen receptors^41^. ERα agonist propyl pyrazole triol (PPT, 250 ng/mouse, Sigma Aldrich, #H6036, St. Louis, MO), or ERβ agonist diarylpropionitrile (DPN, 100 ng/mouse, Sigma Aldrich, #H5915, St. Louis, MO) were subcutaneously injected into ovariectomized mice. Uteri were collected 24 hours after injection. Each group contained at least three mice.

### RNA extraction

Total RNA was extracted using a standard TRIzol reagent (#15596026, Thermo Fisher, Waltham, MA)-based protocol. Briefly, gonads were carefully separated from the mesonephros and then tissues were completely homogenized using TRIzol. Reverse transcription reactions were performed using a LunaScript RT SuperMix Kit (#E3010L, New England Biolabs, Ipswich, MA) with 500 ng of total RNA. The resulting cDNA product was then used for qRT-PCR.

### Quantitative real-time PCR (qRT-PCR)

qRT-PCR was performed on the StepOnePlus Real-Time PCR System (Applied Biosystems, Waltham, MA) using the Fast SYBR Green Master Mix (#4385616, Thermo Fisher, Waltham, MA). Primers were used at a 1 μm concentration, and cycling conditions were 95°C for 20 s, followed by 40 cycles of 95°C for 3 s and 60°C for 30 s. Melt curve analysis or agarose gel electrophoresis was utilized to verify primer specificity. *Gapdh* was used for normalization and all reactions were run in triplicate. Primers used for qRT-PCR are listed in Supplementary Table S1. mRNA expression data from qRT-PCR was calculated relative to controls using the ΔΔCt method. Results were shown as mean ± SD, each with *n*≥3 uteri.

### Primary endometrial stromal cell dissociation and treatment

Endometrial stromal cells were dissociated as previously described^40^. Briefly, uteri were cut open and dissociated in 5mL Hanks’ Balanced Salt Solution (HBSS) containing 6 mg/mL dispase (Thermo Fisher, Waltham, MA) and 1% (w/v) trypsin (Thermo Fisher, Waltham, MA) at 4°C for 45 minutes, room temperature for 20 minutes, and 37°C for 5 minutes to remove luminal epithelial cells. The remaining endometrial tissue pieces were incubated in 5mL HBSS containing 0.15 mg/mL collagenase IV at 37°C for 20 minutes, and the supernatant was filtered through a 70 µm cell strainer to collect stromal cells. Filtered cells were centrifuged at 350 *g* for 10 minutes. Tomato-positive or Venus-positive cells were FACS-purified using a MA900 Multi-Application Cell Sorter (Sony Biotechnology, San Jose, CA).

For DLL4 culture experiments, an 8-well chamber slide (Millipore #PEZGS0816, Burlington, MA) was freshly coated with 2 µg DLL4 (Sino Biological #50640-M08H, Beijing, China) at 37°C for 4 hours. Isolated endometrial cells were seeded into the chamber and cultured with DMEM/F12 containing 10% FBS for 4 days. Cells were treated with 10 nm of estradiol-17β or 10 μM γ-secretase inhibitor IX (DAPT; Calbiochem/EMD Millipore #565770-5MG, Burlington, MA) for 4 days.

### Endometrial organoids

Organoid culture from mouse endometrium was performed as previously described^39^. In brief, uteri were minced into small pieces and dissociated in 1 ml dissociation solution containing 100 µl 12.5 U/ml dispase and 10 µl 40 mg/ml collagen V at 4°C for 30 minutes and 37°C for 10 minutes. The endometrial tissue pieces were mechanically dissociated, and the supernatant was filtered through a 70 µm cell strainer. The filtered cells were centrifuged at 350 g for 10 minutes. The pellet was resuspended in ice-cold DMEM/F12 and centrifuged again. The endometrial cells were placed in 70% Matrigel in media containing a cocktail of growth and signaling factors, which are necessary for mouse endometrial organoid formation^39,79^. To assess the differentiation potential of Nestin+ perivascular cells in organoid assays, Tomato-labeled cells were FACS-purified from P62 *Nestin*-CreER; *Rosa*-Tomato endometrium, exposed to 4-OHT at P60 and P61. Medium was changed every 2 days and organoids were collected after 6 days.

### Single-cell RNA-seq analysis

For Day 4 (D4) analyses, scRNA-seq data of uterine cells on day 4 of pregnancy was downloaded from GEO dataset GSM6213387. Data analysis was conducted using the Seurat package as shown in the published code. After filtering by mitochondrial genes (<50%) and number of gene features (200-8000), 6049 cells were used for subsequent analyses.

For Day 8 (D8) analyses, scRNA-seq data of uterine cells on day 8 of pregnancy was downloaded from GEO dataset GSM5492035. Data analysis was conducted using the Seurat package as shown in the published code. After filtering by mitochondrial genes (<30%) and number of gene features (200-5000), 2349 cells were used for subsequent analyses.

For pericyte analyses, scRNA-seq data of pericytes from *Pdgfrb*-BAC-eGFP mice 24 hours after removal of progesterone was downloaded from GEO dataset GSM5952176. Data analysis was conducted using the Seurat package as shown in the published code. After filtering by mitochondrial genes (<20%) and number of gene features (200-5700), 5059 cells were used for subsequent analyses.

### Immunofluorescence

Whole tissues (uteri) were fixed overnight at 4°C in 4% paraformaldehyde (PFA) with 0.1% Triton X-100. For staining of cell culture samples, cells on slides were fixed in cold 4% PFA with 0.1% Triton X-100 for 15 minutes. After three washes in PBTx (PBS + 0.1% Triton X-100), samples were incubated for 1 hour in blocking solution containing 10% FBS and 3% bovine serum albumin (BSA) at room temperature. Sections were then incubated with primary antibodies (diluted in blocking solution) overnight at 4 °C. After several washes in PBTx, fluorescent secondary antibodies were added for 1 hour at room temperature. Samples were then washed in PBTx; all the remaining liquid was removed; and they were mounted on slides in Fluoromount-G (SouthernBiotech). Primary antibodies and dilutions used in this study are listed in Supplementary Table S2. Nuclear staining was performed using 2 μg/mL Hoechst 33342 (#H1399, Thermo Fisher, Waltham, MA), and Hoechst staining is labeled in figures as “Nuclei”. Secondary antibodies used in this study include Alexa-488-, Alexa-555-, and Alexa-647-conjugated secondary antibodies (Life Technologies, Carlsbad, CA), used at 1:500 dilution. Pictures were taken on a Nikon A1 inverted LUNV microscope (Nikon, Tokyo, Japan) equipped with Nikon’s NIS-Elements imaging software (Nikon, Tokyo, Japan). At least two independent experiments were performed and within each experiment multiple uteri (*n*≥3) were used.

### Statistical analysis

All statistical analyses were performed using Prism version 8.0 (GraphPad). Two-tailed Student t-tests were performed to calculate *P* values, with *P*<0.05 considered statistically significant. Error bars indicate +/- SEM. For immunofluorescence, organoid, and qRT-PCR assays, at least three independent experiments were performed, and within each experiment multiple uteri (*n*≥3) were used. The mouse model used and the time points for sample collection in each experiment are described in individual figure legends. ImageJ was used for cell counting and co-localization analysis. The JACoP Plugin within ImageJ was applied to evaluate co-localization. Each dot on graphs represents an individual, independent measurement from one mouse; for every experimental condition, at least three mice (*n*≥3) were used.

---

**Supplementary Figure 1.**
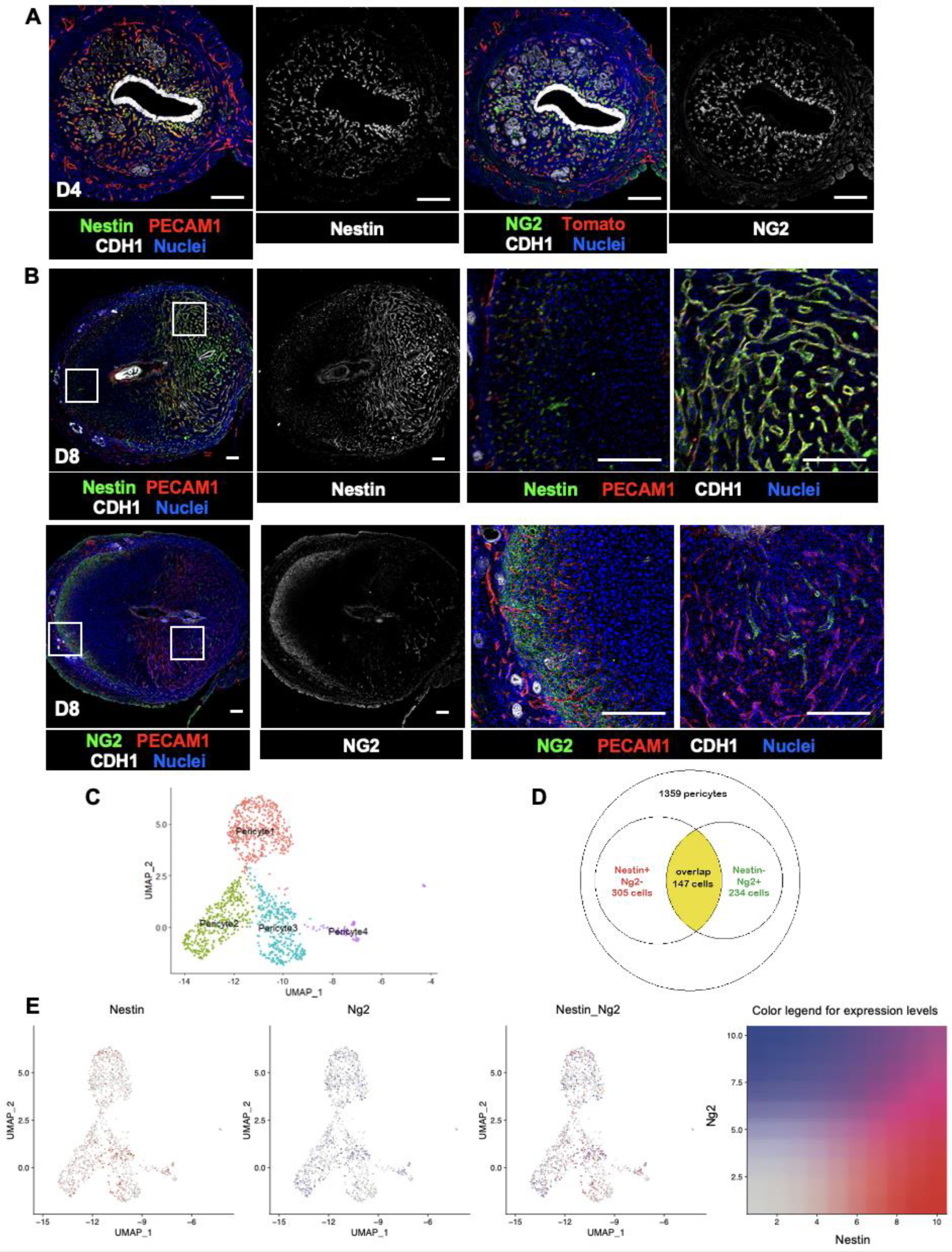
Dynamic expression pattern of Nestin and NG2 during uterine regeneration. (A, B) Immunofluorescence images of D4 (A) and D8 (B) uteri. Scale bar: 200 μm. (C) UMAP visualization of pericyte subpopulations from a previously published scRNA-seq dataset^16^. (D) Venn diagram showing quantification of *Nestin*+ and/or *Ng2*+ pericytes among all perivascular cells from scRNA-seq data. (E) Feature plots showing expression profiles of *Nestin* (left), *Ng2* (middle) and the overlap of both (right) in perivascular populations. Color codes for expression levels are presented in the color gradient image.

**Supplementary Figure 2.**
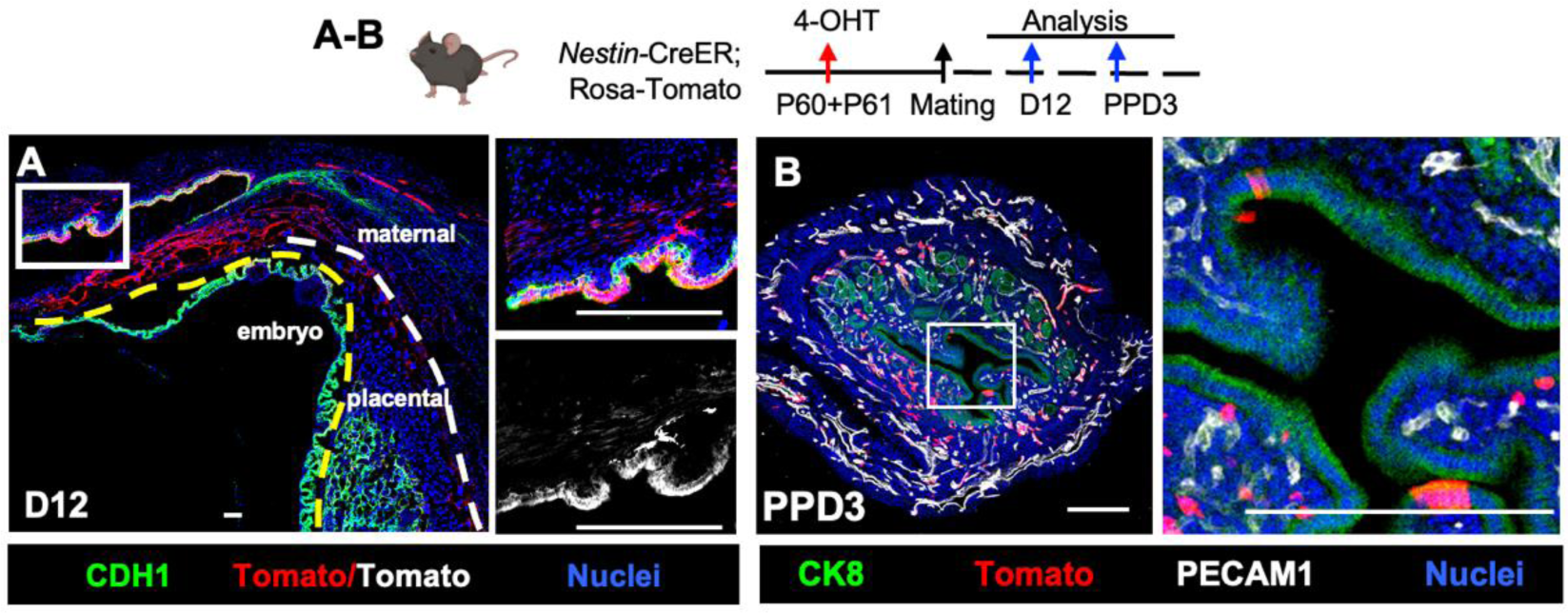
The contribution of *Nestin*+ cells to epithelial regeneration during pregnancy. (A, B) Immunofluorescence images of *Nestin*-CreER; *Rosa*-Tomato uteri at D12 (A) and PPD3 (B) following exposure to 4-OHT at P60 and P61. Scale bar: 100 μm.

**Supplementary Figure 3.**
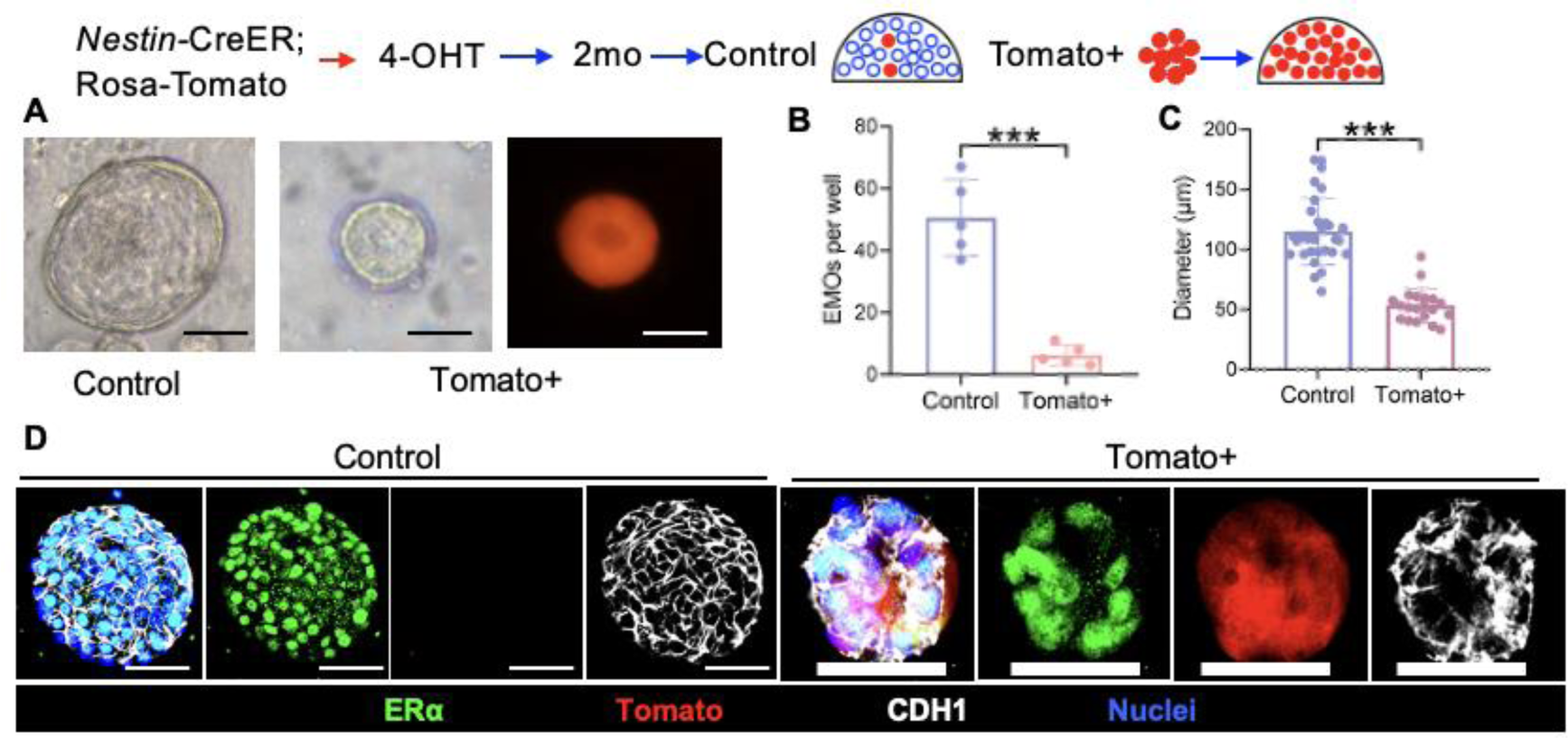
Endometrial organoids can be generated using Tomato+ cells from P62 *Nestin*-CreER; *Rosa*-Tomato endometrium. (A) Representative images of organoids from whole endometrial cells (control) and FACS-purified Tomato+ cells alone. (B-C) Quantification of number (B) and diameter (C) of organoids. (D) Immunofluorescence images showing that organoids expressed ERα and the epithelial marker CDH1. No Tomato+ cells were observed in control (whole tissue) organoids. Scale bar: 50 μm. \*\*\**P*<0.001.

**Supplementary Figure 4.**
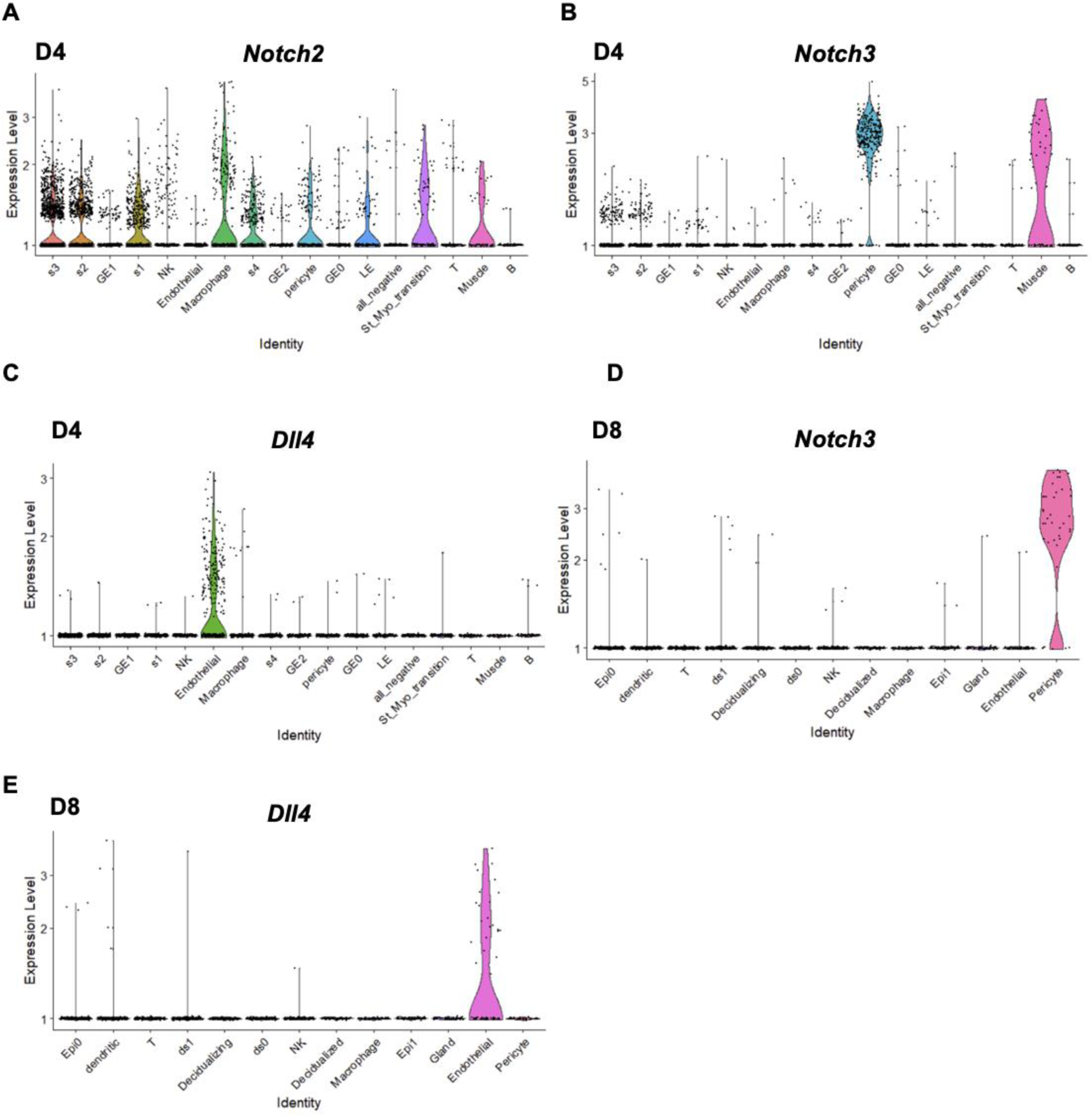
The cellular distribution of Notch and its ligands in the pregnant mouse uterus. (A-E) Violin plots from previously published uterine scRNA-seq data showing the distribution and expression levels of *Notch2* (A), *Notch3* (B), and *Dll4* (C) in different cell types in D4 mouse uterus^43^, and *Notch3* (D) and *Dll4* (E) in different cell types in D8 mouse uterus ^44^.

**Supplementary Figure 5.**
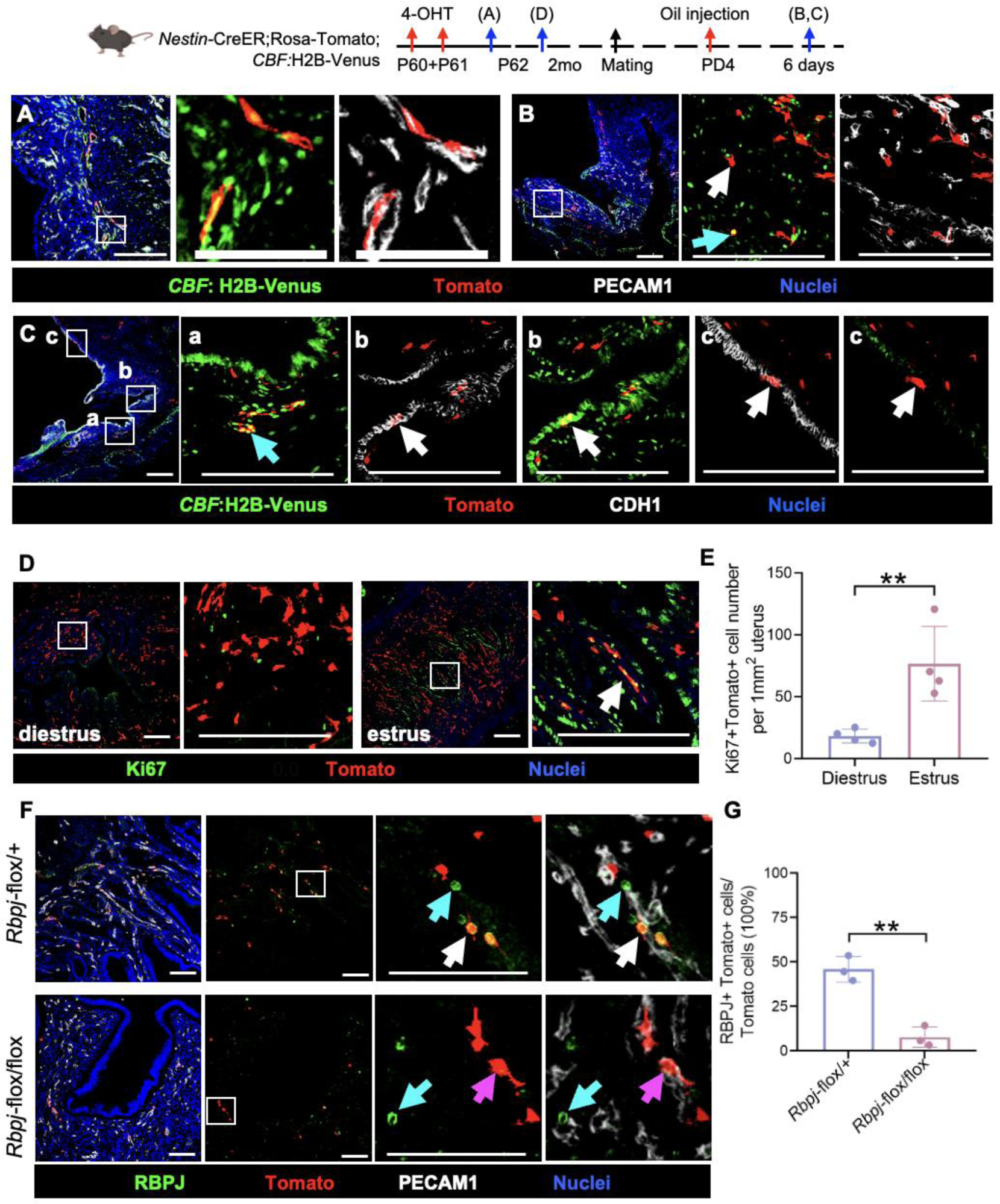
*Nestin*+ perivascular cells exhibit active Notch signaling and are in the quiescent phase of the cell cycle. (A-D) Immunofluorescence images of *Nestin*-CreER; *Rosa*-Tomato; *CBF*:H2B-Venus uteri at various time points after exposure to 4-OHT at P60 and P61 (see timeline in schematic). White arrow in B indicates perivascular Tomato cell, while cyan arrow points to Venus-expressing Tomato+ cell. Cyan arrow in C-a indicates perivascular Venus+ Tomato cell, and white arrows point to epithelial Tomato+ cells with (C-b) and without (C-c) Venus expression. (E) Quantification of Ki67+ Tomato+ cell number per unit area. (F) Immunofluorescence images of *Nestin*-CreER; *Rosa*-Tomato; *Rpbj*-flox/+ heterozygous and *Nestin*-CreER; *Rosa*-Tomato; *Rpbj*-flox/flox uteri 2 days after exposure to 4-OHT at the diestrus phase. White arrow indicates RBPJ+ Tomato+ cell, while magenta arrow points to RBPJ-negative Tomato+ cell. Cyan arrow points to Tomato-negative perivascular cell that expresses RBPJ. (G) Quantification of percent Tomato+ cells expressing RBPJ. Scale bar: 100 μm. \*\**P*<0.01.

**Supplementary Table S1.**
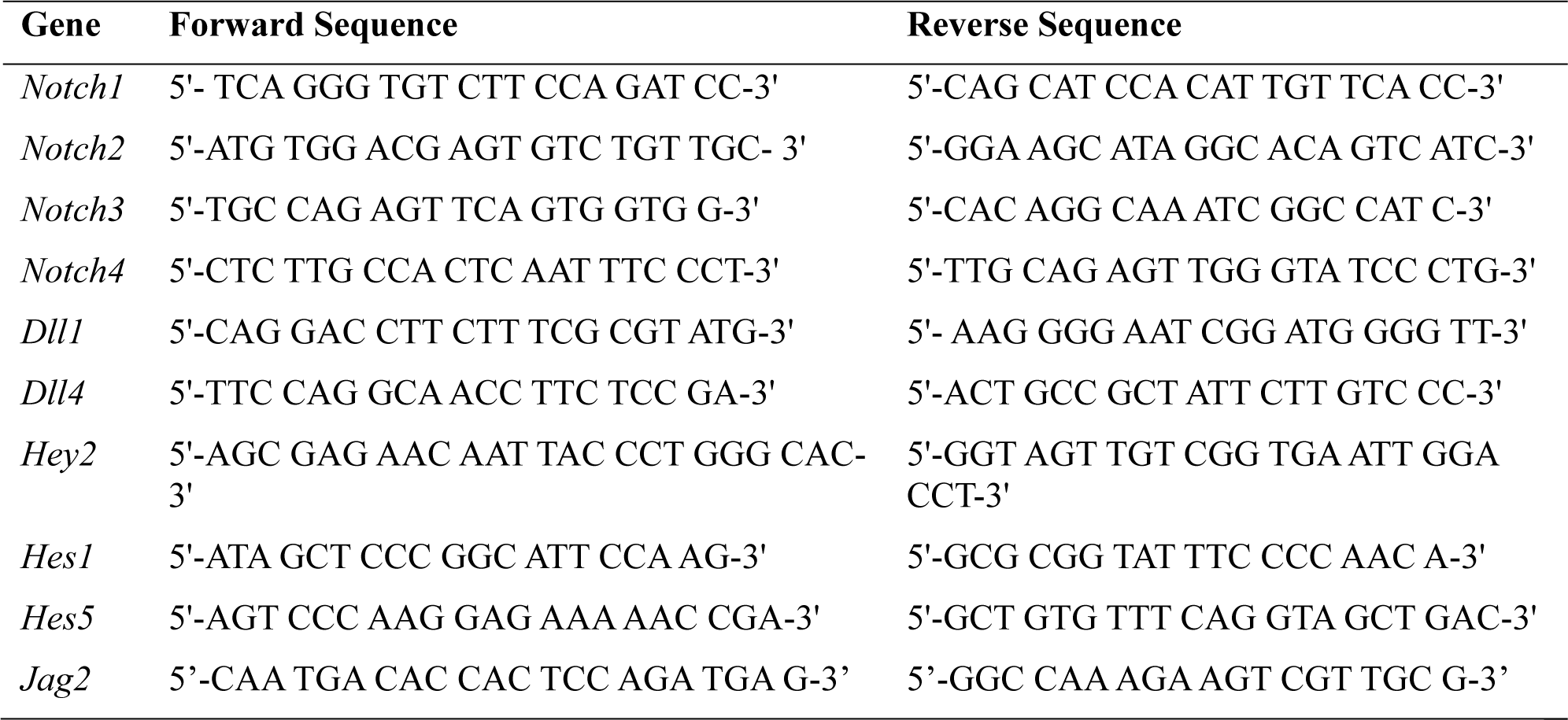
List of primer sequences used for qRT-PCR.

**Supplementary Table S2.** List of primary antibodies used.

| Primary Antibody | Dilution | Source |
| --- | --- | --- |
| Rabbit anti-Nestin | 1:1000 | BioLegend #PRB-315C |
| Rat anti-PECAM1 | 1:250 | BD Pharmingen #553370 |
| Rabbit anti-ERG | 1:250 | Abcam #ab92513 |
| Rat anti-CK8 | 1:500 | EMD Milipore #MABT329 |
| Rabbit anti-NG2 | 1:500 | Millipore-Sigma #4147 |
| Rat anti-CDH1 | 1:500 | Invitrogen #13-1900 |
| Rabbit anti-ERα | 1:200 | Santa Cruz #sc-542 |
| Rabbit anti-NOTCH3 | 1:1000 | Cell Signaling #5276T |
| Rabbit anti-Ki67 | 1:300 | Thermo RM-9106-S |
| Rabbit anti-RBPJ | 1:1000 | Cell Signaling Technology #5314 |
| Rat anti-Ki67 | 1:500 | Thermo 14-5698-80 |
| Goat anti-DLL4 | 1:100 | R&D Systems #AF389 |

## Notes

### Competing Interest Statement

The authors have declared no competing interest.

## References

1. Figueira, P.G., Abrao, M.S., Krikun, G., and Taylor, H.S. (2011). Stem cells in endometrium and their role in the pathogenesis of endometriosis. Ann N Y Acad Sci 1221, 10–17. 10.1111/j.1749-6632.2011.05969.x.

2. Cooke, P.S., Spencer, T.E., Bartol, F.F., and Hayashi, K. (2013). Uterine glands: development, function and experimental model systems. Mol Hum Reprod 19, 547–558. 10.1093/molehr/gat031.

3. Gargett, C.E., Nguyen, H.P., and Ye, L. (2012). Endometrial regeneration and endometrial stem/progenitor cells. Rev Endocr Metab Disord 13, 235–251. 10.1007/s11154-012-9221-9.

4. Ng, S.W., Norwitz, G.A., Pavlicev, M., Tilburgs, T., Simon, C., and Norwitz, E.R. (2020). Endometrial Decidualization: The Primary Driver of Pregnancy Health. Int J Mol Sci 21. 10.3390/ijms21114092.

5. Critchley, H.O.D., Maybin, J.A., Armstrong, G.M., and Williams, A.R.W. (2020). Physiology of the Endometrium and Regulation of Menstruation. Physiol Rev 100, 1149–1179. 10.1152/physrev.00031.2019.

6. Jin, S. (2019). Bipotent stem cells support the cyclical regeneration of endometrial epithelium of the murine uterus. Proc Natl Acad Sci U S A 116, 6848–6857. 10.1073/pnas.1814597116.

7. Sanchez-Mata, A., and Gonzalez-Munoz, E. (2021). Understanding menstrual blood-derived stromal/stem cells: Definition and properties. Are we rushing into their therapeutic applications? iScience 24, 103501. 10.1016/j.isci.2021.103501.

8. Salamonsen, L.A., Hutchison, J.C., and Gargett, C.E. (2021). Cyclical endometrial repair and regeneration. Development 148. 10.1242/dev.199577.

9. Li, S., and Ding, L. (2021). Endometrial Perivascular Progenitor Cells and Uterus Regeneration. J Pers Med 11. 10.3390/jpm11060477.

10. Cousins, F.L., Murray, A., Esnal, A., Gibson, D.A., Critchley, H.O., and Saunders, P.T. (2014). Evidence from a mouse model that epithelial cell migration and mesenchymal-epithelial transition contribute to rapid restoration of uterine tissue integrity during menstruation. PLoS One 9, e86378. 10.1371/journal.pone.0086378.

11. Syed, S.M., Kumar, M., Ghosh, A., Tomasetig, F., Ali, A., Whan, R.M., Alterman, D., and Tanwar, P.S. (2020). Endometrial Axin2(+) Cells Drive Epithelial Homeostasis, Regeneration, and Cancer following Oncogenic Transformation. Cell Stem Cell 26, 64–80 e13. 10.1016/j.stem.2019.11.012.

12. Gao, Y., Li, S., and Li, Q. (2014). Uterine epithelial cell proliferation and endometrial hyperplasia: evidence from a mouse model. Mol Hum Reprod 20, 776–786. 10.1093/molehr/gau033.

13. Zhai, J., Vannuccini, S., Petraglia, F., and Giudice, L.C. (2020). Adenomyosis: Mechanisms and Pathogenesis. Semin Reprod Med 38, 129–143. 10.1055/s-0040-1716687.

14. Huang, C.C., Orvis, G.D., Wang, Y., and Behringer, R.R. (2012). Stromal-to-epithelial transition during postpartum endometrial regeneration. PLoS One 7, e44285. 10.1371/journal.pone.0044285.

15. Patterson, A.L., Zhang, L., Arango, N.A., Teixeira, J., and Pru, J.K. (2013). Mesenchymal-to-epithelial transition contributes to endometrial regeneration following natural and artificial decidualization. Stem Cells Dev 22, 964–974. 10.1089/scd.2012.0435.

16. Kirkwood, P.M., Gibson, D.A., Shaw, I., Dobie, R., Kelepouri, O., Henderson, N.C., and Saunders, P.T.K. (2022). Single-cell RNA sequencing and lineage tracing confirm mesenchyme to epithelial transformation (MET) contributes to repair of the endometrium at menstruation. Elife 11. 10.7554/eLife.77663.

17. Crisan, M., Corselli, M., Chen, C.W., and Peault, B. (2011). Multilineage stem cells in the adult: a perivascular legacy? Organogenesis 7, 101–104. 10.4161/org.7.2.16150.

18. Cao, D., Chan, R.W.S., Ng, E.H.Y., Gemzell-Danielsson, K., and Yeung, W.S.B. (2021). Single-cell RNA sequencing of cultured human endometrial CD140b(+)CD146(+) perivascular cells highlights the importance of in vivo microenvironment. Stem Cell Res Ther 12, 306. 10.1186/s13287-021-02354-1.

19. Zhou, B., Lin, W., Long, Y., Yang, Y., Zhang, H., Wu, K., and Chu, Q. (2022). Notch signaling pathway: architecture, disease, and therapeutics. Signal Transduct Target Ther 7, 95. 10.1038/s41392-022-00934-y.

20. Siebel, C., and Lendahl, U. (2017). Notch Signaling in Development, Tissue Homeostasis, and Disease. Physiol Rev 97, 1235–1294. 10.1152/physrev.00005.2017.

21. Massri, N., Loia, R., Sones, J.L., Arora, R., and Douglas, N.C. (2023). Vascular changes in the cycling and early pregnant uterus. JCI Insight 8. 10.1172/jci.insight.163422.

22. Cuman, C., Menkhorst, E., Winship, A., Van Sinderen, M., Osianlis, T., Rombauts, L.J., and Dimitriadis, E. (2014). Fetal-maternal communication: the role of Notch signalling in embryo implantation. Reproduction 147, R75–86. 10.1530/REP-13-0474.

23. Reichrath, J., and Reichrath, S. (2020). Notch Signaling and Embryonic Development: An Ancient Friend, Revisited. Adv Exp Med Biol 1218, 9–37. 10.1007/978-3-030-34436-8_2.

24. Afshar, Y., Jeong, J.W., Roqueiro, D., DeMayo, F., Lydon, J., Radtke, F., Radnor, R., Miele, L., and Fazleabas, A. (2012). Notch1 mediates uterine stromal differentiation and is critical for complete decidualization in the mouse. FASEB J 26, 282–294. 10.1096/fj.11-184663.

25. Zhang, S., Chan, R.W.S., Ng, E.H.Y., and Yeung, W.S.B. (2022). The role of Notch signaling in endometrial mesenchymal stromal/stem-like cells maintenance. Commun Biol 5, 1064. 10.1038/s42003-022-04044-x.

26. Zhang, S., Kong, S., Wang, B., Cheng, X., Chen, Y., Wu, W., Wang, Q., Shi, J., Zhang, Y., Wang, S., et al. (2014). Uterine Rbpj is required for embryonic-uterine orientation and decidual remodeling via Notch pathway-independent and -dependent mechanisms. Cell Res 24, 925–942. 10.1038/cr.2014.82.

27. Wiese, C., Rolletschek, A., Kania, G., Blyszczuk, P., Tarasov, K.V., Tarasova, Y., Wersto, R.P., Boheler, K.R., and Wobus, A.M. (2004). Nestin expression--a property of multi-lineage progenitor cells? Cell Mol Life Sci 61, 2510–2522. 10.1007/s00018-004-4144-6.

28. Bernal, A., and Arranz, L. (2018). Nestin-expressing progenitor cells: function, identity and therapeutic implications. Cell Mol Life Sci 75, 2177–2195. 10.1007/s00018-018-2794-z.

29. Carlen, M., Meletis, K., Barnabe-Heider, F., and Frisen, J. (2006). Genetic visualization of neurogenesis. Exp Cell Res 312, 2851–2859. 10.1016/j.yexcr.2006.05.012.

30. Imayoshi, I., Ohtsuka, T., Metzger, D., Chambon, P., and Kageyama, R. (2006). Temporal regulation of Cre recombinase activity in neural stem cells. Genesis 44, 233–238. 10.1002/dvg.20212.

31. Kuo, C.T., Mirzadeh, Z., Soriano-Navarro, M., Rasin, M., Wang, D., Shen, J., Sestan, N., Garcia-Verdugo, J., Alvarez-Buylla, A., Jan, L.Y., and Jan, Y.N. (2006). Postnatal deletion of Numb/Numblike reveals repair and remodeling capacity in the subventricular neurogenic niche. Cell 127, 1253–1264. 10.1016/j.cell.2006.10.041.

32. Battiste, J., Helms, A.W., Kim, E.J., Savage, T.K., Lagace, D.C., Mandyam, C.D., Eisch, A.J., Miyoshi, G., and Johnson, J.E. (2007). Ascl1 defines sequentially generated lineage-restricted neuronal and oligodendrocyte precursor cells in the spinal cord. Development 134, 285–293. 10.1242/dev.02727.

33. Burns, K.A., Ayoub, A.E., Breunig, J.J., Adhami, F., Weng, W.L., Colbert, M.C., Rakic, P., and Kuan, C.Y. (2007). Nestin-CreER mice reveal DNA synthesis by nonapoptotic neurons following cerebral ischemia hypoxia. Cereb Cortex 17, 2585–2592. 10.1093/cercor/bhl164.

34. Cicero, S.A., Johnson, D., Reyntjens, S., Frase, S., Connell, S., Chow, L.M., Baker, S.J., Sorrentino, B.P., and Dyer, M.A. (2009). Cells previously identified as retinal stem cells are pigmented ciliary epithelial cells. Proc Natl Acad Sci U S A 106, 6685–6690. 10.1073/pnas.0901596106.

35. Li, S.Y., Bhandary, B., Gu, X., and DeFalco, T. (2022). Perivascular cells support folliculogenesis in the developing ovary. Proc Natl Acad Sci U S A 119, e2213026119. 10.1073/pnas.2213026119.

36. Kumar, D.L., and DeFalco, T. (2018). A perivascular niche for multipotent progenitors in the fetal testis. Nat Commun 9, 4519. 10.1038/s41467-018-06996-3.

37. Cousins, F.L., Pandoy, R., Jin, S., and Gargett, C.E. (2021). The Elusive Endometrial Epithelial Stem/Progenitor Cells. Front Cell Dev Biol 9, 640319. 10.3389/fcell.2021.640319.

38. Li, S.Y., Song, Z., Song, M.J., Qin, J.W., Zhao, M.L., and Yang, Z.M. (2016). Impaired receptivity and decidualization in DHEA-induced PCOS mice. Sci Rep 6, 38134. 10.1038/srep38134.

39. Seishima, R., Leung, C., Yada, S., Murad, K.B.A., Tan, L.T., Hajamohideen, A., Tan, S.H., Itoh, H., Murakami, K., Ishida, Y., et al. (2019). Neonatal Wnt-dependent Lgr5 positive stem cells are essential for uterine gland development. Nat Commun 10, 5378. 10.1038/s41467-019-13363-3.

40. Li, S.Y., Yan, J.Q., Song, Z., Liu, Y.F., Song, M.J., Qin, J.W., Yang, Z.M., and Liang, X.H. (2017). Molecular characterization of lysyl oxidase-mediated extracellular matrix remodeling during mouse decidualization. FEBS Lett 591, 1394–1407. 10.1002/1873-3468.12645.

41. Liu, Y.F., Li, M.Y., Yan, Y.P., Wei, W., Li, B., Pan, H.Y., Yang, Z.M., and Liang, X.H. (2020). ERalpha-dependent stimulation of LCN2 in uterine epithelium during mouse early pregnancy. Reproduction 159, 493–501. 10.1530/REP-19-0616.

42. Nowotschin, S., Xenopoulos, P., Schrode, N., and Hadjantonakis, A.K. (2013). A bright single-cell resolution live imaging reporter of Notch signaling in the mouse. BMC Dev Biol 13, 15. 10.1186/1471-213X-13-15.

43. Li, R., Wang, T., Marquardt, R.M., Lydon, J.P., Wu, S.P., and DeMayo, F.J. (2023). TRIM28 modulates nuclear receptor signaling to regulate uterine function. Nat Commun 14, 4605. 10.1038/s41467-023-40395-7.

44. Li, R., Wang, T.Y., Xu, X., Emery, O.M., Yi, M., Wu, S.P., and DeMayo, F.J. (2022). Spatial transcriptomic profiles of mouse uterine microenvironments at pregnancy day 7.5dagger. Biol Reprod 107, 529–545. 10.1093/biolre/ioac061.

45. Urban, N., and Cheung, T.H. (2021). Stem cell quiescence: the challenging path to activation. Development 148. 10.1242/dev.165084.

46. Bjornson, C.R., Cheung, T.H., Liu, L., Tripathi, P.V., Steeper, K.M., and Rando, T.A. (2012). Notch signaling is necessary to maintain quiescence in adult muscle stem cells. Stem Cells 30, 232–242. 10.1002/stem.773.

47. Cheung, T.H., and Rando, T.A. (2013). Molecular regulation of stem cell quiescence. Nat Rev Mol Cell Biol 14, 329–340. 10.1038/nrm3591.

48. Uxa, S., Castillo-Binder, P., Kohler, R., Stangner, K., Muller, G.A., and Engeland, K. (2021). Ki-67 gene expression. Cell Death Differ 28, 3357–3370. 10.1038/s41418-021-00823-x.

49. Zakrzewski, W., Dobrzynski, M., Szymonowicz, M., and Rybak, Z. (2019). Stem cells: past, present, and future. Stem Cell Res Ther 10, 68. 10.1186/s13287-019-1165-5.

50. Pittenger, M.F., Discher, D.E., Peault, B.M., Phinney, D.G., Hare, J.M., and Caplan, A.I. (2019). Mesenchymal stem cell perspective: cell biology to clinical progress. NPJ Regen Med 4, 22. 10.1038/s41536-019-0083-6.

51. Zhao, X., and Moore, D.L. (2018). Neural stem cells: developmental mechanisms and disease modeling. Cell Tissue Res 371, 1–6. 10.1007/s00441-017-2738-1.

52. Tan, S., and Barker, N. (2015). Epithelial stem cells and intestinal cancer. Semin Cancer Biol 32, 40–53. 10.1016/j.semcancer.2014.02.005.

53. Dzierzak, E., and Bigas, A. (2018). Blood Development: Hematopoietic Stem Cell Dependence and Independence. Cell Stem Cell 22, 639–651. 10.1016/j.stem.2018.04.015.

54. Hong, I.S. (2023). Endometrial stem/progenitor cells: Properties, origins, and functions. Genes Dis 10, 931–947. 10.1016/j.gendis.2022.08.009.

55. Ferenczy, A. (1976). Studies on the cytodynamics of human endometrial regeneration. I. Scanning electron microscopy. Am J Obstet Gynecol 124, 64–74. 10.1016/0002-9378(76)90013-2.

56. Valentijn, A.J., Palial, K., Al-Lamee, H., Tempest, N., Drury, J., Von Zglinicki, T., Saretzki, G., Murray, P., Gargett, C.E., and Hapangama, D.K. (2013). SSEA-1 isolates human endometrial basal glandular epithelial cells: phenotypic and functional characterization and implications in the pathogenesis of endometriosis. Hum Reprod 28, 2695–2708. 10.1093/humrep/det285.

57. Ali, A., Syed, S.M., Jamaluddin, M.F.B., Colino-Sanguino, Y., Gallego-Ortega, D., and Tanwar, P.S. (2020). Cell Lineage Tracing Identifies Hormone-Regulated and Wnt-Responsive Vaginal Epithelial Stem Cells. Cell Rep 30, 1463–1477 e1467. 10.1016/j.celrep.2020.01.003.

58. Spitzer, T.L., Rojas, A., Zelenko, Z., Aghajanova, L., Erikson, D.W., Barragan, F., Meyer, M., Tamaresis, J.S., Hamilton, A.E., Irwin, J.C., and Giudice, L.C. (2012). Perivascular human endometrial mesenchymal stem cells express pathways relevant to self-renewal, lineage specification, and functional phenotype. Biol Reprod 86, 58. 10.1095/biolreprod.111.095885.

59. Crisan, M., Corselli, M., Chen, W.C., and Peault, B. (2012). Perivascular cells for regenerative medicine. J Cell Mol Med 16, 2851–2860. 10.1111/j.1582-4934.2012.01617.x.

60. Mannino, G., Russo, C., Maugeri, G., Musumeci, G., Vicario, N., Tibullo, D., Giuffrida, R., Parenti, R., and Lo Furno, D. (2022). Adult stem cell niches for tissue homeostasis. J Cell Physiol 237, 239–257. 10.1002/jcp.30562.

61. Oh, M., and Nor, J.E. (2015). The Perivascular Niche and Self-Renewal of Stem Cells. Front Physiol 6, 367. 10.3389/fphys.2015.00367.

62. Li, S., Bhandary, B., and DeFalco, T. (2021). Perivascular cells support folliculogenesis in the developing ovary. 2021.2004.2026.441453. 10.1101/2021.04.26.441453 %J bioRxiv.

63. Akil, A., Gutierrez-Garcia, A.K., Guenter, R., Rose, J.B., Beck, A.W., Chen, H., and Ren, B. (2021). Notch Signaling in Vascular Endothelial Cells, Angiogenesis, and Tumor Progression: An Update and Prospective. Front Cell Dev Biol 9, 642352. 10.3389/fcell.2021.642352.

64. Imayoshi, I., Sakamoto, M., Yamaguchi, M., Mori, K., and Kageyama, R. (2010). Essential roles of Notch signaling in maintenance of neural stem cells in developing and adult brains. J Neurosci 30, 3489–3498. 10.1523/JNEUROSCI.4987-09.2010.

65. Foster, J.S., Henley, D.C., Ahamed, S., and Wimalasena, J. (2001). Estrogens and cell-cycle regulation in breast cancer. Trends Endocrinol Metab 12, 320–327. 10.1016/s1043-2760(01)00436-2.

66. Chan, R.W., Kaitu’u-Lino, T., and Gargett, C.E. (2012). Role of label-retaining cells in estrogen-induced endometrial regeneration. Reprod Sci 19, 102–114. 10.1177/1933719111414207.

67. Cooke, P.S., Buchanan, D.L., Young, P., Setiawan, T., Brody, J., Korach, K.S., Taylor, J., Lubahn, D.B., and Cunha, G.R. (1997). Stromal estrogen receptors mediate mitogenic effects of estradiol on uterine epithelium. Proc Natl Acad Sci U S A 94, 6535–6540. 10.1073/pnas.94.12.6535.

68. Pawar, S., Laws, M.J., Bagchi, I.C., and Bagchi, M.K. (2015). Uterine Epithelial Estrogen Receptor-alpha Controls Decidualization via a Paracrine Mechanism. Mol Endocrinol 29, 1362–1374. 10.1210/me.2015-1142.

69. Winuthayanon, W., Lierz, S.L., Delarosa, K.C., Sampels, S.R., Donoghue, L.J., Hewitt, S.C., and Korach, K.S. (2017). Juxtacrine Activity of Estrogen Receptor alpha in Uterine Stromal Cells is Necessary for Estrogen-Induced Epithelial Cell Proliferation. Sci Rep 7, 8377. 10.1038/s41598-017-07728-1.

70. Kaitu’u-Lino, T.J., Morison, N.B., and Salamonsen, L.A. (2007). Estrogen is not essential for full endometrial restoration after breakdown: lessons from a mouse model. Endocrinology 148, 5105–5111. 10.1210/en.2007-0716.

71. Gkikas, D., Tsampoula, M., and Politis, P.K. (2017). Nuclear receptors in neural stem/progenitor cell homeostasis. Cell Mol Life Sci 74, 4097–4120. 10.1007/s00018-017-2571-4.

72. Mao, A., Paharkova-Vatchkova, V., Hardy, J., Miller, M.M., and Kovats, S. (2005). Estrogen selectively promotes the differentiation of dendritic cells with characteristics of Langerhans cells. J Immunol 175, 5146–5151. 10.4049/jimmunol.175.8.5146.

73. Okazaki, R., Inoue, D., Shibata, M., Saika, M., Kido, S., Ooka, H., Tomiyama, H., Sakamoto, Y., and Matsumoto, T. (2002). Estrogen promotes early osteoblast differentiation and inhibits adipocyte differentiation in mouse bone marrow stromal cell lines that express estrogen receptor (ER) alpha or beta. Endocrinology 143, 2349–2356. 10.1210/endo.143.6.8854.

74. Gong, H., Chen, Y., Xu, J., Xie, X., Yu, D., Yang, B., and Kuang, H. (2017). The regulation of ovary and conceptus on the uterine natural killer cells during early pregnancy. Reprod Biol Endocrinol 15, 73. 10.1186/s12958-017-0290-1.

75. Borzychowski, A.M., Chantakru, S., Minhas, K., Paffaro, V.A., Yamada, A.T., He, H., Korach, K.S., and Croy, B.A. (2003). Functional analysis of murine uterine natural killer cells genetically devoid of oestrogen receptors. Placenta 24, 403–411. 10.1053/plac.2002.0924.

76. Tanigaki, K., Han, H., Yamamoto, N., Tashiro, K., Ikegawa, M., Kuroda, K., Suzuki, A., Nakano, T., and Honjo, T. (2002). Notch-RBP-J signaling is involved in cell fate determination of marginal zone B cells. Nat Immunol 3, 443–450. 10.1038/ni793.

77. Sano, K., Matsuda, S., Tohyama, S., Komura, D., Shimizu, E., and Sutoh, C. (2020). Deep learning-based classification of the mouse estrous cycle stages. Sci Rep 10, 11714. 10.1038/s41598-020-68611-0.

78. Byers, S.L., Wiles, M.V., Dunn, S.L., and Taft, R.A. (2012). Mouse estrous cycle identification tool and images. PLoS One 7, e35538. 10.1371/journal.pone.0035538.

79. Gu, Z.Y., Jia, S.Z., Liu, S., and Leng, J.H. (2020). Endometrial Organoids: A New Model for the Research of Endometrial-Related Diseasesdagger. Biol Reprod 103, 918–926. 10.1093/biolre/ioaa124.

